# Major axes of variation in tree demography across global forests

**DOI:** 10.1101/2023.01.11.523538

**Authors:** Melina de Souza Leite, Sean M. McMahon, Paulo Inácio Prado, Stuart J. Davies, Alexandre Adalardo de Oliveira, Hannes P. De Deurwaerder, Salomón Aguilar, Kristina J. Anderson-Teixeira, Nurfarah Aqilah, Norman A. Bourg, Warren Y. Brockelman, Nicolas Castaño, Chia-Hao Chang-Yang, Yu-Yun Chen, George Chuyong, Keith Clay, Álvaro Duque, Sisira Ediriweera, Corneille E.N. Ewango, Gregory Gilbert, I.A.U.N. Gunatilleke, C.V.S. Gunatilleke, Robert Howe, Walter Huaraca Huasco, Akira Itoh, Daniel J. Johnson, David Kenfack, Kamil Král, Yao Tze Leong, James A. Lutz, Jean-Remy Makana, Yadvinder Malhi, William J. McShea, Mohizah Mohamad, Musalmah Nasardin, Anuttara Nathalang, Geoffrey Parker, Renan Parmigiani, Rolando Pérez, Richard P. Phillips, Pavel Šamonil, I-Fang Sun, Sylvester Tan, Duncan Thomas, Jill Thompson, María Uriarte, Amy Wolf, Jess Zimmerman, Daniel Zuleta, Marco D. Visser, Lisa Hülsmann

## Abstract

The future trajectory of global forests is closely intertwined with tree demography, and a major fundamental goal in ecology is to understand the key mechanisms governing spatial-temporal patterns in tree population dynamics. While historical research has made substantial progress in identifying the mechanisms individually, their relative importance among forests remains unclear mainly due to practical limitations. One approach is to group mechanisms according to their shared effects on the variability of tree vital rates and to quantify patterns therein. We developed a conceptual and statistical framework (variance partitioning of Bayesian multilevel models) that attributes the variability in tree growth, mortality, and recruitment to variation in species, space, and time, and their interactions, categories we refer to as *organising principles* (OPs). We applied the framework to data from 21 forest plots covering more than 2.9 million trees of approximately 6,500 species. We found that differences among species, the *species* OP, proved a major source of variability in tree vital rates, explaining 28-33% of demographic variance alone, and in interaction with *space* 14-17%, totalling 40-43%. The average variability among species declined with species richness across forests, indicating that diverse forests featured smaller interspecific differences in vital rates supporting the theory that the range of vital rates is similar across global forests. Decomposing the variance in vital rates into the proposed OPs showed that taxonomy is crucial to predicting and understanding tree demography on large forest plots. A focus on how variance is organized in forests can facilitate the construction of more targeted models with clearer expectations of which covariates might drive a vital rate. This study therefore highlights the most promising avenues for future research, both in terms of understanding the relative contributions of groups of mechanisms to forest demography and diversity, and for improving projections of forest ecosystems.

## Introduction

Forests are an integral component of the global carbon cycle (Anderson-Teixeira *et al*., 2021) and are home to a majority of the terrestrial biodiversity (Pillay *et al*., 2022). Changes in climate and land use threaten forests but anticipating how these diverse systems might respond is challenged by the broad array of mechanisms that might determine forest structure and function. Further, these mechanisms differ in their influence over space and time and are difficult to measure at the appropriate scale of their potential influence. A common approach to quantifying forest function is through the analysis of tree demography (Griffith *et al*., 2016): the growth, survival, and reproduction of individual trees. These vital (i.e. demographic) rates combine to determine key features of forests, such as biomass stocks and fluxes (Needham *et al*., 2022), structural complexity (Kohyama, 1993), and diversity (Lasky *et al*., 2014). A better understanding of forest demography can advance the development and testing of ecological theories such as the role of coexistence (Broekman *et al*., 2019; Hülsmann *et al*., 2020) and niche (Kohyama, 1993; Lasky *et al*., 2014) in community ecology. Moreover, demography has been identified as critical for more accurately modelling the terrestrial component in earth system models (Fisher *et al*., 2018) and projecting the future of the terrestrial carbon sink (Pan *et al*., 2011). Even small changes, over space and time, in tree vital rates can affect carbon cycles (Needham *et al*., 2022) and thus the extent to which climate change can be mitigated by forests (Canadell & Raupach, 2008).

Vital rates are influenced by interacting mechanisms across spatial and temporal scales creating a challenge to the inclusion of demography in forest models (Weng *et al*., 2015). Many of these mechanisms are difficult or impossible to measure directly, leading to the use of imperfect proxies (Swenson *et al*., 2020). Besides, data analysis is usually restricted to a few non-interacting proxies, making even harder to distinguish and compare mechanisms’ relative importance. There exists, however, a higher level of information that may guide demographic analyses focused on mechanisms: the patterns in vital rates themselves. The three vital rates and the contextual variables (‘dimensions’) associated with them offer an opportunity to organise the elements of forest dynamics in ways that help to infer the potential mechanisms that structure forests. For example, through natural selection species have developed different strategies to acquire and allocate resources. This results in a species dimension that represents the range of phenotypes among species (Díaz *et al*., 2016) and, thus, also the observed vital rates of individual species (Johnson *et al*., 2018; Rüger *et al*., 2018; Needham *et al*., 2022). Moreover, as resource availability and stressors vary along spatial and temporal dimensions, the environmental conditions of a forest also structure the vital rates of the trees, e.g. soil and topography vary across space (Zuleta *et al*., 2020) and drought conditions over time (Chen *et al*., 2019). Finally, all these dimensions (species, space, and time) have interactive effects. Functional traits vary between species and cause differential responses along spatial and temporal dimensions, for example when drought tolerant and intolerant species respond differently to a climatic event (Kupers *et al*., 2019). Gap dynamics change over both space and time, and tree responses change as forest gaps close (Wright *et al*., 2003). Patterns of how variability in vital rates is partitioned along these key dimensions can thus reveal how important various biotic and abiotic drivers are in influencing tree demography and by extension forest dynamics.

We present a conceptual framework that groups the mechanisms creating variation in vital rates as being related to species, space, and time. Together, these three dimensions and their interactions form seven organising principles (OPs, Table 1). When the mechanisms that drive tree vital rates operate on unique combinations of these dimensions, quantifying the variability in vital rates that each OP describes may will provide insights into the strength and relative importance of the mechanisms that might potentially be correlated with that rate (Table 1). The statistical counterpart to this conceptual framework is variance partitioning analysis, a technique that decomposes the variability in the response variable to the groups of interest in the data (Searle *et al*., 2006), using multilevel models (McMahon & Diez, 2007; Visser *et al*., 2016). In our framework, we decompose forest demographic data across OPs and quantify the relative importance of each OP by estimating and partitioning the variance in each vital rate (Browne *et al*., 2005). By attributing the total variability in vital rates to the different OPs, a broad assessment of the structure of variation in vital rates can be accomplished (Table 1).

**Table 1.**
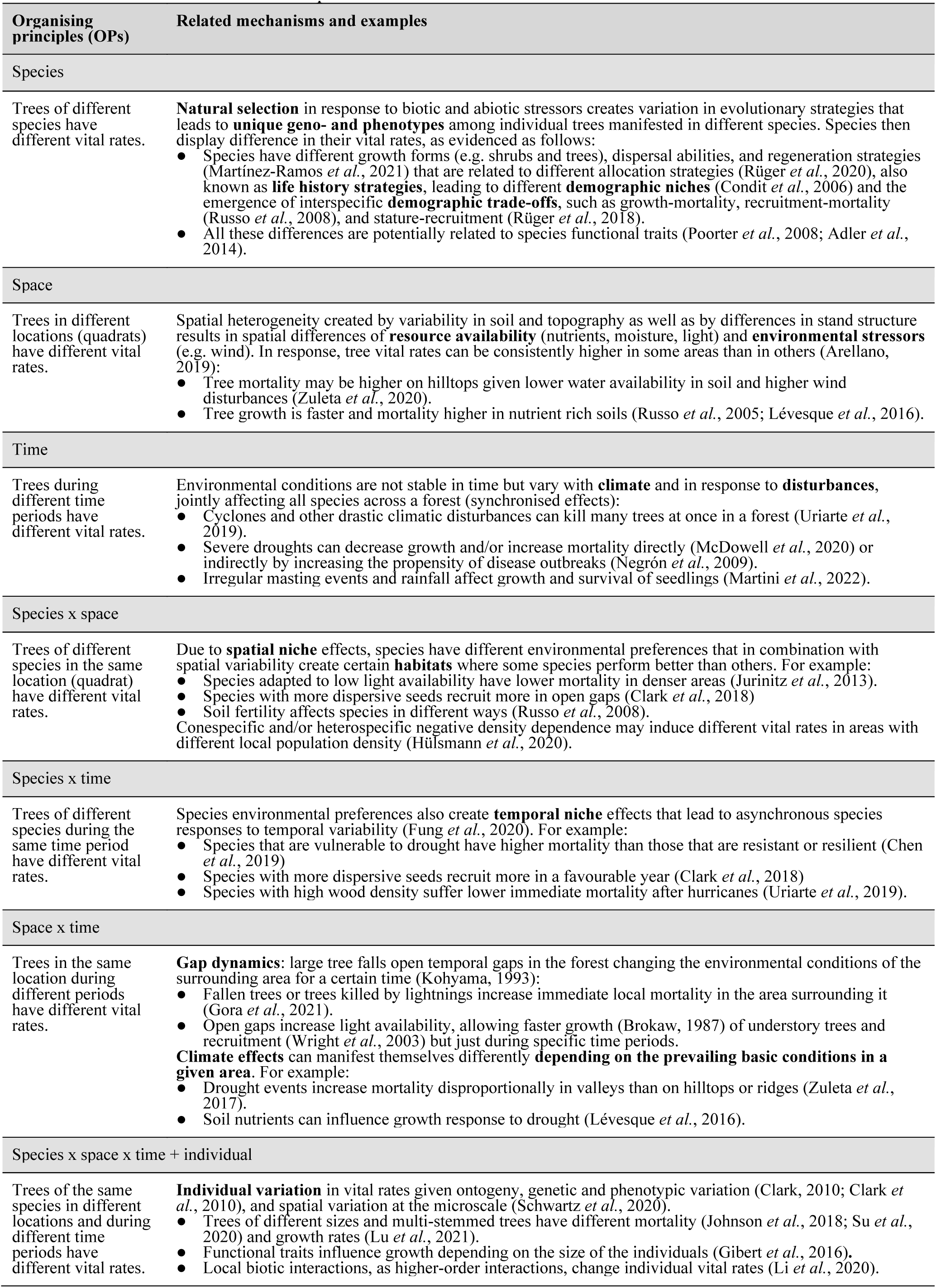
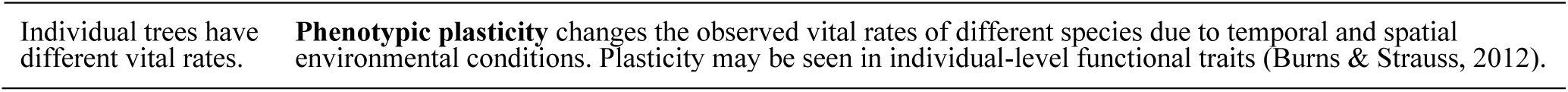
Seven organising principles (OPs) and the mechanisms that are associated with them, i.e., by creating variability of vital rates in the associated dimensions *species, space* and *time* and their interactions. References are example studies for the mechanisms.

| Organising principles (OPs) | Related mechanisms and examples |
| --- | --- |
| **Species** |  |
| Trees of different species have different vital rates. | **Natural selection** in response to biotic and abiotic stressors creates variation in evolutionary strategies that leads to **unique geno- and phenotypes** among individual trees manifested in different species. Species then display difference in their vital rates, as evidenced as follows:<br>Species have different growth forms (e.g. shrubs and trees), dispersal abilities, and regeneration strategies (Martínez-Ramos *et al.*, 2021) that are related to different allocation strategies (Rüger *et al.*, 2020), also known as **life history strategies**, leading to different **demographic niches** (Condit *et al.*, 2006) and the emergence of interspecific **demographic trade-offs**, such as growth-mortality, recruitment-mortality (Russo *et al.*, 2008), and stature-recruitment (Rüger *et al.*, 2018).<br>All these differences are potentially related to species functional traits (Poorter *et al.*, 2008; Adler *et al.*, 2014). |
| **Space** |  |
| Trees in different locations (quadrats) have different vital rates. | Spatial heterogeneity created by variability in soil and topography as well as by differences in stand structure results in spatial differences of **resource availability** (nutrients, moisture, light) and **environmental stressors** (e.g. wind). In response, tree vital rates can be consistently higher in some areas than in others (Arellano, 2019):<br>Tree mortality may be higher on hilltops given lower water availability in soil and higher wind disturbances (Zuleta *et al.*, 2020).<br>Tree growth is faster and mortality higher in nutrient rich soils (Russo *et al.*, 2005; Lévesque *et al.*, 2016). |
| **Time** |  |
| Trees during different time periods have different vital rates. | Environmental conditions are not stable in time but vary with **climate** and in response to **disturbances**, jointly affecting all species across a forest (synchronised effects):<br>Cyclones and other drastic climatic disturbances can kill many trees at once in a forest (Uriarte *et al.*, 2019).<br>Severe droughts can decrease growth and/or increase mortality directly (McDowell *et al.*, 2020) or indirectly by increasing the propensity of disease outbreaks (Negrón *et al.*, 2009).<br>Irregular masting events and rainfall affect growth and survival of seedlings (Martini *et al.*, 2022). |
| **Species x space** |  |
| Trees of different species in the same location (quadrat) have different vital rates. | Due to **spatial niche** effects, species have different environmental preferences that in combination with spatial variability create certain **habitats** where some species perform better than others. For example:<br>Species adapted to low light availability have lower mortality in denser areas (Jurinitz *et al.*, 2013).<br>Species with more dispersive seeds recruit more in open gaps (Clark *et al.*, 2018)<br>Soil fertility affects species in different ways (Russo *et al.*, 2008).<br>Conespecific and/or heterospecific negative density dependence may induce different vital rates in areas with different local population density (Hülsmann *et al.*, 2020). |
| **Species x time** |  |
| Trees of different species during the same time period have different vital rates. | Species environmental preferences also create **temporal niche** effects that lead to asynchronous species responses to temporal variability (Fung *et al.*, 2020). For example:<br>Species that are vulnerable to drought have higher mortality than those that are resistant or resilient (Chen *et al.*, 2019)<br>Species with more dispersive seeds recruit more in a favourable year (Clark *et al.*, 2018)<br>Species with high wood density suffer lower immediate mortality after hurricanes (Uriarte *et al.*, 2019). |
| **Space x time** |  |
| Trees in the same location during different periods have different vital rates. | **Gap dynamics:** large tree falls open temporal gaps in the forest changing the environmental conditions of the surrounding area for a certain time (Kohyama, 1993):<br>Fallen trees or trees killed by lightnings increase immediate local mortality in the area surrounding it (Gora *et al.*, 2021).<br>Open gaps increase light availability, allowing faster growth (Brokaw, 1987) of understory trees and recruitment (Wright *et al.*, 2003) but just during specific time periods.<br>**Climate effects** can manifest themselves differently **depending on the prevailing basic conditions in a given area**. For example:<br>Drought events increase mortality disproportionally in valleys than on hilltops or ridges (Zuleta *et al.*, 2017).<br>Soil nutrients can influence growth response to drought (Lévesque *et al.*, 2016). |
| **Species x space x time + individual** |  |
| Trees of the same species in different locations and during different time periods have different vital rates. | **Individual variation** in vital rates given ontogeny, genetic and phenotypic variation (Clark, 2010; Clark *et al.*, 2010), and spatial variation at the microscale (Schwartz *et al.*, 2020).<br>Trees of different sizes and multi-stemmed trees have different mortality (Johnson *et al.*, 2018; Su *et al.*, 2020) and growth rates (Lu *et al.*, 2021).<br>Functional traits influence growth depending on the size of the individuals (Gibert *et al.*, 2016).<br>Local biotic interactions, as higher-order interactions, change individual vital rates (Li *et al.*, 2020). |

We applied this framework to a set of 21 large (6 to 52 ha) and globally distributed forest dynamics plots (Davies *et al*., 2021). We then compared the relative importance of the OPs for each vital rate at each forest with the goal of identifying consistent patterns in which OPs capture variation in vital rates: (1) among vital rates, i.e. investigating if some OPs are more important than others for specific vital rates; (2) across spatial scales (grain size), given the nature of scale dependency of ecological processes; and (3) among forests globally to understand how patterns may differ depending on forest diversity and structure. In doing so, we provide macroecological patterns of the relative importance of OPs and, thus, the first approximate assessment of their associated mechanisms in generating variation in global forest demography. Our framework aims to facilitate hypothesis-driven research on mechanisms by first describing the higher-level patterns of vital rate variability and giving important insights to the ecological dimensions ‘at which the action lies’ (Browne et al. 2005).

## Methods

### Tree census data

We used data from 21 forest dynamics plots (Fig 1A) from the Forest Global Earth Observatory network (ForestGEO, Davies *et al*., 2021). In each plot, all stems with dbh ≥1 cm were mapped, identified, and repeatedly measured using a standardised protocol. Plots used in this study range in size between 6 and 52 ha, with an inter-census measurement interval of approximately 5 years (range 3 to 10 years). The area within each forest plot was subdivided into quadrats of equal sizes (see *Organizing principles across spatial scales*). All forest plots had at least 2 censuses. The forest plots cover a wide range of environmental, climatic, and edaphic conditions, with the number of species per plot varying two orders of magnitude from 12 to 1402 (including morphospecies). In total, approximately 2.9 million trees from more than 6,500 species were repeatedly censused over periods of 3 to 40 years in more than 575 ha. For summary information on the plots and further details on how tree census data were processed see Appendix S1 in Supporting Information.

**Figure 1.**
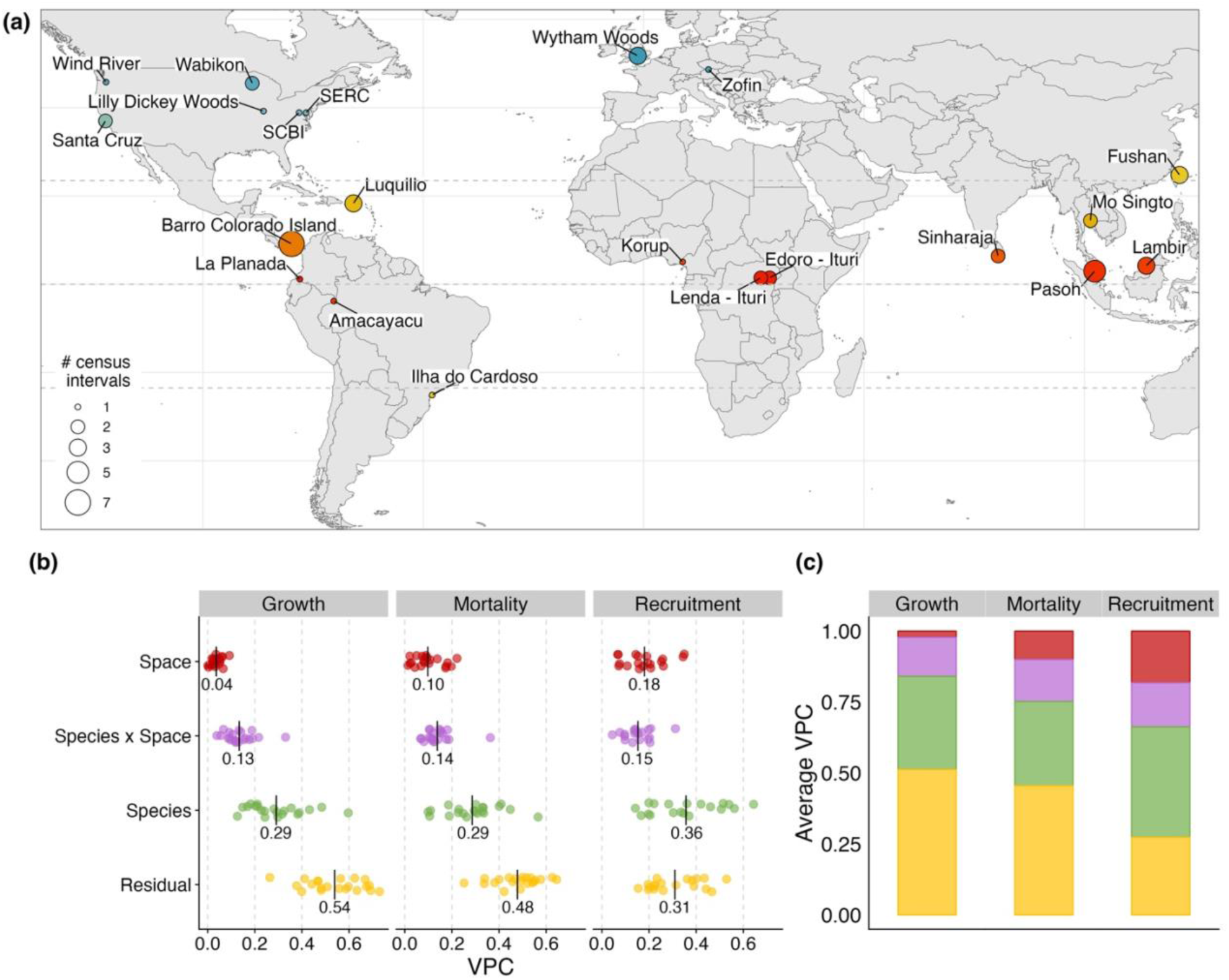
(a) Global distribution of the 21 forest plots. (b) Variance partition coefficients (VPC) of the organising principles (OPs) per vital rate - growth, mortality, and recruitment - with mean values indicated as black vertical lines and numbers. (c) Average VPCs across all plots, where colours correspond to the same OPs as in (b). Models were fitted at the 5x5 m grain size. Each forest plot in (a) is coloured by latitude and the size of the circle is related to the number of census intervals.

### Vital rate definition and modelling

We analysed growth, mortality, and recruitment as annual rates by using vital rate information at the level of individual trees and applying the variance partitioning analysis per forest plot and vital rate. Annual individual growth was calculated as dbh increment in millimetres of alive trees, divided by the individual’s census interval length in years, and modelled using multilevel models with a normal distribution.

Variance partitioning of mortality and recruitment is less intuitive than growth, because although every individual has a unique, observable growth rate, individual trees only provide an observable status (i.e., individuals are either alive or dead). However, we can estimate latent mortality and recruitment rates for individuals belonging to the same population, space, and time by calculating per-capita vital rates (sensu Kohyama *et al*., 2018). Further, although the variance of individual binary observations is fixed at 1.68 (the standard deviation of a logistic distribution [see below]), this term has meaning when compared to other sources of measurable variance, such as across populations, years, or spatial aggregations. Therefore, mortality was estimated from the status of trees - alive or dead - in each consecutive census assuming a binomial distribution (Kohyama *et al*., 2018). Mortality rates were annualised by using a complementary log log link function (cloglog), where the log-transformed time between individual measurements is included as an offset term (Fortin *et al*., 2008; Johnson *et al*., 2018).

Recruitment was defined as the final per-capita recruitment rate (Kohyama *et al*., 2018), which denotes the proportion of trees that are new recruits (i.e. not present in the previous census) and can be interpreted as the probability of an individual tree being new. Recruitment rates were estimated using the same modelling approach as for mortality, i.e., a binomial model with a cloglog link function and time interval length as an offset term. Because there is no time interval associated with individual recruits as they have not been monitored in the previous census, the time interval for recruitment was calculated as the mean time interval of the survivors in the same quadrat. If there were no survivors in a specific quadrat, we used the mean time interval between the respective censuses from the entire plot.

### Variance partitioning analysis

To quantify the variation in vital rates associated with each OP, we applied variance partitioning to multilevel models (MLMs) fitted separately for each vital rate and forest plot. MLMs are particularly useful for variance decomposition as they are able to reflect that ecological datasets contain identifiable hierarchical units, groups, or clusters (McMahon & Diez, 2007). MLMs can account for such interdependence by partitioning the total variance into different components of variation due to each cluster (see example in Table 1). We included *species*, quadrat (*space*) and census interval (*time*) and their two-way interactions as variance components. With that, we estimated the variance associated with each OP while respecting the hierarchical structure of the data. Following the convention of MLMs, the general structure of our models is:

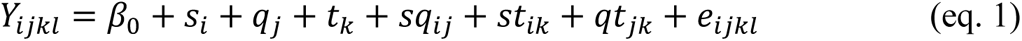

where 𝑌_𝑖𝑗𝑘𝑙_ is the vital rate for individual observation *l*, from species *i*, in quadrat *j* and time interval *k*. 𝛽_0_ is the global intercept. 𝑠_𝑖_, 𝑞_𝑗_, 𝑡_𝑘_ are effects at the species, space (quadrat) and time level, respectively (commonly termed as random effects in mixed-effects models), while 𝑠𝑞_𝑖𝑗_, 𝑠𝑡_𝑖𝑘_, 𝑞𝑡_𝑗𝑘_ are effects of interactions between OPs: *species x space*, *species x time*, and *space x time*. All parameters are represented by a normal distribution with mean zero and their respective variances 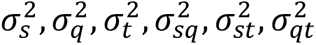 with no covariance being modelled. The residual variance (𝑒_𝑖𝑗𝑘𝑙_) represents the variance explained by the three-way interaction *species x space x time*, and any unexplained variation among observations including non-separable measurement error and individual variation (Table 1). Residual variance in growth models assume a normally distributed error. For mortality and recruitment, modelled with binomial distributions, the residual variance at the link scale (i.e. linear predictor scale) is the expected variance for the binomial distribution (𝜋^2^/6 ∼1.68) (Nakagawa *et al*., 2017). We decided not to include the three-way interaction *species x space x time* because most of the clusters formed by the combinations of the *species*, *space*, and *time* categories would have only one tree, i.e., not enough observations per cluster especially for the small spatial grains, preventing the model to correctly compute variance among these clusters. The same reasoning applies to the individual variance, where the number of individual trees with only one measurement is high for all forests plots. It means that any variability given to the tree-way interaction *species x space x tim*e and the individual variation will be attributed to the residual variance in growth models (normal distribution) and will not be accounted for in the mortality and recruitment models (binomial distribution.

To partition the total variance of the vital rates among the individual OPs, we calculated **variance partition coefficients (VPCs)** (Browne *et al*., 2005). The VPC of each OP was calculated as the proportion of its variance to the total variance of the model. It is worth noting that we intentionally included no fixed effects in the models, in contrast to the usual statistical approach when searching for specific mechanisms, because all mechanisms are considered through OPs, which represent the dimensions at which they generate variability.

All data analyses were performed using R (R Core Team, 2022), using the R package ‘brms‘ (Bürkner, 2017) to build Bayesian MLMs. For all estimated parameters, we used brms default weakly informative prior distributions. For each model, we ran three Monte Carlo Markov chains with 3,000 iterations, discarding the first 1,000 iterations and thinning with an interval of 5, resulting in 1200 posterior samples. We checked convergence of the chains using the Gelman– Rubin criterion and by visually inspecting trace plots of estimated coefficients.

### Analysis framework

#### Organising principles among vital rates

To assess the relative importance of the OPs among vital rates, we compared the VPC results for each vital rate among the 21 forest plots. However, because 16 forests had too few census intervals (see below), i.e., less than three (Table S1.2), we fit a reduced version of the variance partitioning analysis (eq. 1) without the temporal OPs (dropping the variances 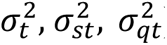). We ran separate analysis for each time interval of the same forest plot and averaged variances for forests with more than one census interval.

#### Temporal organising principles

Currently, a bottleneck of our analysis is the scarcity of data for the temporal dimension of vital rates variability. For variance partitioning analysis, the estimation of the variance of a grouping variable (i.e., *time* OP) with less than four to five levels may be biased towards zero (Oberpriller *et al*., 2022). In our data, only five forest plots in the (sub)tropics (Table S1.2) presented a reasonable number of census intervals (i.e., at least four censuses spanning between 20 and 40 years) to be considered suitable for the VPC analysis including temporal OPs (eq.1). We fit these MLMs to ten random subsets of 5 ha each sampled from the full forest plots, where each subset was composed of five non-overlapping quadrats of 1 ha. This procedure was necessary to restrict computational time resulting from the large number of observations, especially on the large plots that are species-rich and of high tree density (i.e., Barro Colorado Island 50 ha, Lambir 50 ha, Pasoh 50 ha, Fig 1a and Table S1.2). Variance estimates of the OPs for each forest plot were averaged across estimates of the ten subsets.

#### Organising principles across spatial scales

To assess how the relative importance of OP varies with spatial scale, i.e., how the choice of a specific grain size impacts VPCs, we divided each forest plot into non-overlapping quadrats with increasing size: 5x5 m (0.0025 ha), 10x10 m (0.01 ha), 20x20 m (0.04 ha), 50x50 m (0.25 ha) and 100x100 m (1 ha). Depending on the size of the plot, we trimmed the data to fit within a rectangular region with edges that were even multiples of 100 m, discarding the data outside this area. This guaranteed that each plot could be evenly divided into quadrats of 1 ha and that the same area was analysed at all spatial scales. We ran variance partitioning analyses without and with temporal OPs averaged VPCs over all forest plots for each grain size and vital rate.

#### Organising principles across a global species richness gradient

Globally, species richness is one of the most distinguishing characteristics of forests and strongly correlates, for instance, with latitude (Keil & Chase, 2019), precipitation (Adler & Levine, 2007), and biome history (Wiens & Donoghue, 2004). The plots used in this analysis span two orders of magnitude in the number of species (12 to 1402, including morphospecies) offering a unique opportunity to explore if and how sources of variability in vital rates are associated with species diversity. We therefore assessed how log-transformed rarefied species richness (cf. Appendix S5) is associated with the VPCs of *species*, *space*, *species x space* and *residual* OP using dirichlet regression from the R package ‘DirichletReg‘ (Maier, 2021), which is appropriate for response variables that are multiple categories of proportional data (Douma & Weedon, 2019).

### Robustness analyses

We performed four extra analyses to make sure our VPCs estimates from the forest plots are robust to (1) different forest plot sizes (6 to 50 ha) for the models without temporal OPs, by subsampling and comparing VPCs of the same forest (Lambir) with the entire plot data (Appendix S2); (2) to the approach of computing average VPCs for the model with temporal OPs from subsampled plots (10 samples of 5 ha each) (Appendix S2); (3) to changes in the modelling procedure, by including or excluding temporal OPs from the VPC analysis (Appendix S3); and (4) to the presence of rare species on VPCs by excluding or including rare species (Appendix S4).

VPCs estimates from all forest plots were robust to changes in plot size and subsampling data. VPC estimates also remained reliable after removing temporal OPs. Specifically, our main results are also robust to the presence of rare species, though excluding or regrouping rare species does result in small decreases in the *species* VPC, balanced by an increase in the *residual* and *species x space* VPC (Appendix S4).

## Results

### Organising principles among vital rates

When comparing the relative importance of the OPs for all 21 forests, we found that, despite large differences among the plots with respect to climate, environment, species richness etc., the relative importance of the OPs was relatively similar (Fig. 1). Generally, *species* was the most important OP for explaining variance in all three vital rates, after the *residual* OP. At the smallest spatial grain (quadrats at 5x5 m), average *species* and *species x space* VPCs varied little among vital rates, ranging from 29 to 36%, and 13 to 15%, respectively. The average *space* VPC was smaller for growth (4%), intermediate for mortality (10%) and larger for recruitment (19%). *Residual* VPCs were on average about half of the total variance for growth and mortality (55 and 47%, respectively) but smaller for recruitment (31%).

### Temporal organising principles

When analysing demographic data from the five forest plots with more than four consecutive censuses (grain size 5x5 m), we found that *species* remained the most important OP to explain variance in tree vital rates, except for growth, where the *species x space* VPC was larger for four of the five plots (Fig. 2). Temporal OPs (*time*, *species x time* and *space x time*) were especially important for mortality and recruitment, where VPCs of *space x time* (on average 10 and 15%, respectively) were larger than VPCs of *species x space* (on average 6 and 10%, respectively).

**Figure 2.**
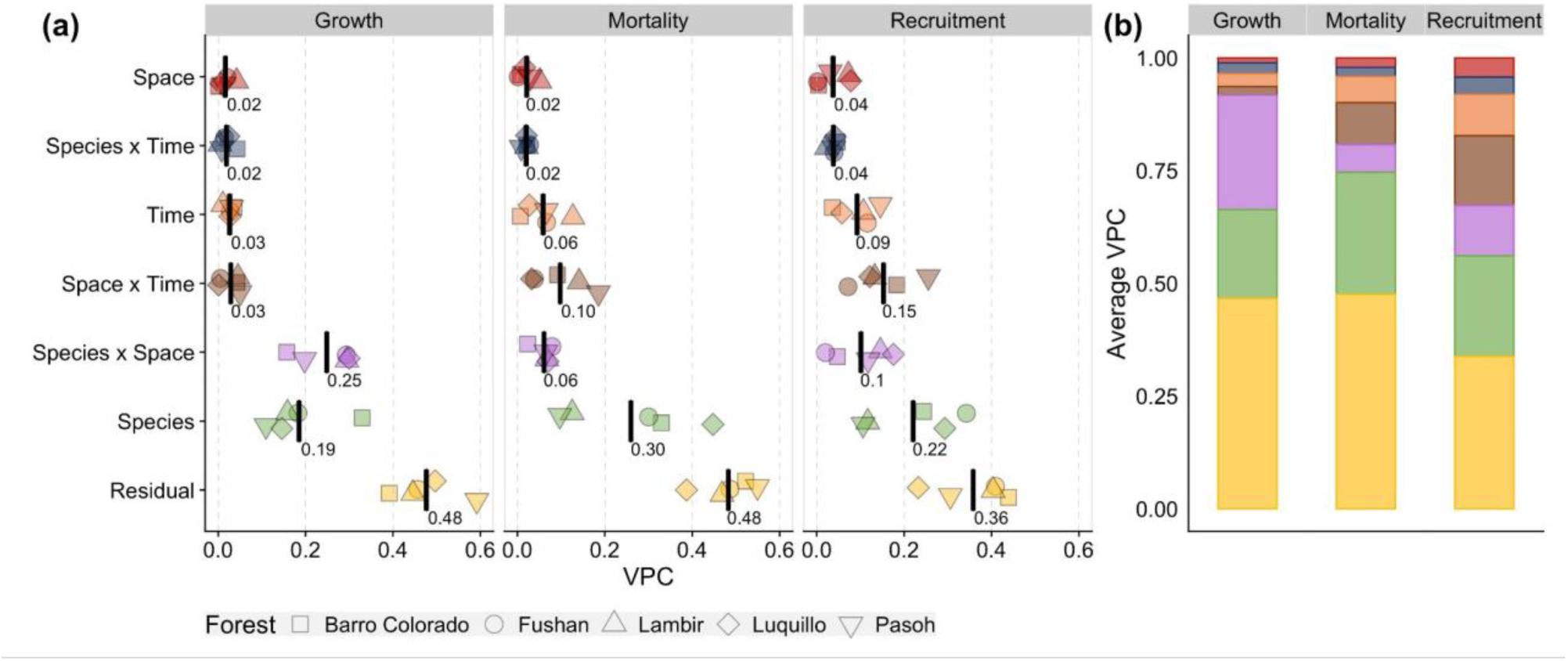
(a) Variance partition coefficient (VPC) of the organising principles (OPs) per vital rate - growth, mortality, and recruitment - for the five forest plots with at least four censuses (different point shapes). Average VPCs across plots are presented as black lines and numbers. (b) Average VPCs across the five plots, where colours correspond to the same OPs as in (a). Models were fitted at the 5x5 m grain size. See Fig. 1a for forest plot locations.

### Organising principles across spatial scales

When comparing average VPCs across five spatial grain sizes, we found that the relative importance of residual variation increased with grain size for all vital rates and more accentuated for growth (Fig. 3). For instance, for the models including temporal OPs (Fig. 3b), residual variation increased from 46% at the smallest grain (quadrats at 5x5 m) to 71% at the largest grain (100x100 m). In turn, the spatial OPs - *space*, *species x space* and *space x time* - consistently decreased in relative importance with increasing spatial grain for all vital rates. The OPs *species* and *species x time* remained almost equally important across spatial grains.

**Figure 3.**
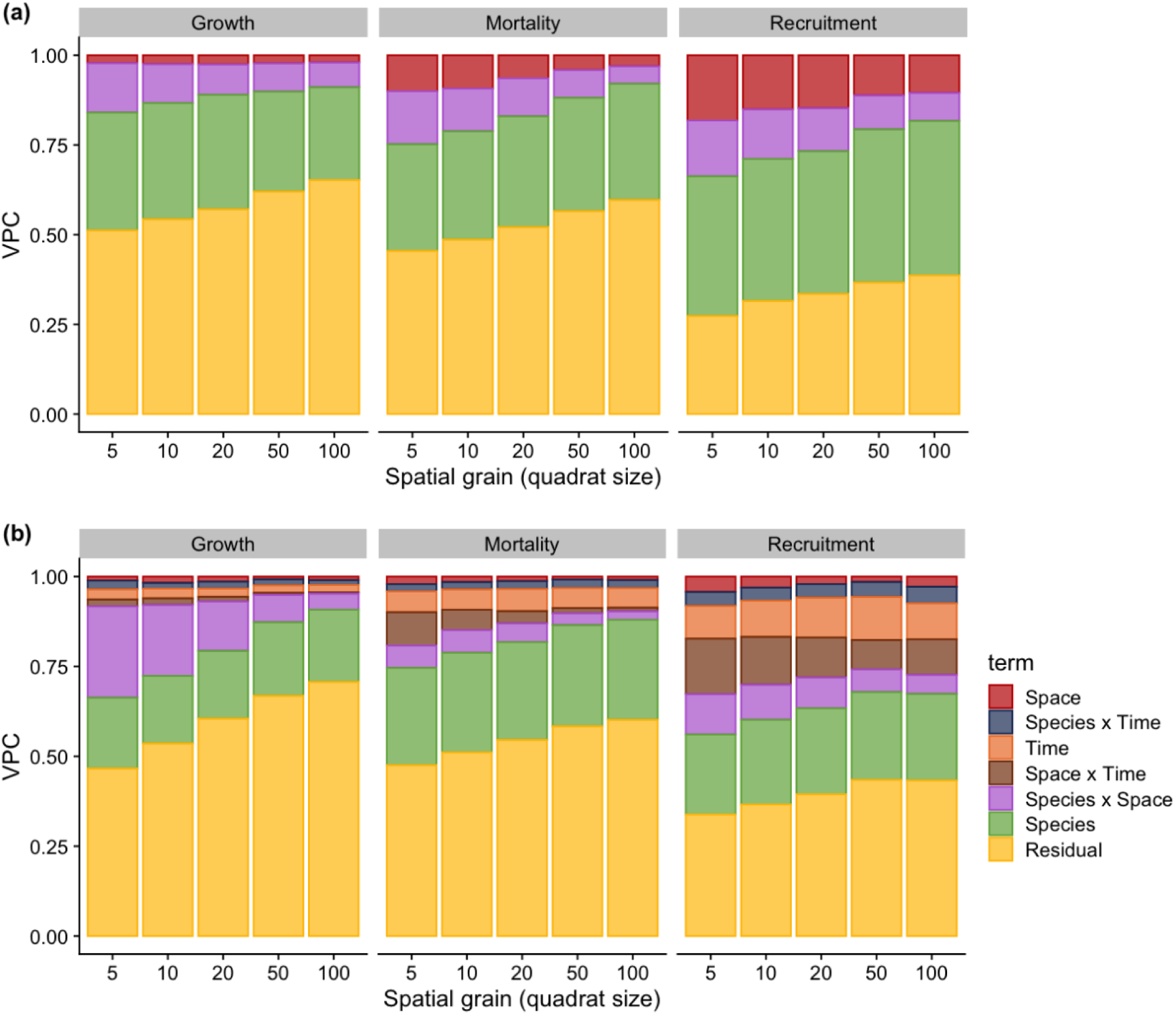
Average variance partition coefficients (VPCs) of each organising principle (OP) across five spatial grain sizes (quadrats from 5x5 m to 100x100 m) for the vital rates growth, mortality, and recruitment: (a) reduced models without temporal OPs for all 21 forests plots, and (b) full models with temporal OPs for the five (sub)tropical forest plots with enough censuses (Barro Colorado Island, Fushan, Lambir, Luquillo, and Pasoh).

### Organising principles across a global species richness gradient

While the *species* OP was the most important VPC for the vital rates throughout the forests, we also found that the importance of the *species* OP decreased with species richness for recruitmet and growth, but not for mortality (Fig. 4). The decrease in the *species* VPC for growth and recruitment was led by a decrease in the *species* standard deviation (Fig. S5.4). This result was robust to the presence of rare species (Fig. S5.5). The other OPs showed no significant changes with species richness (Fig. 4).

**Figure 4.**
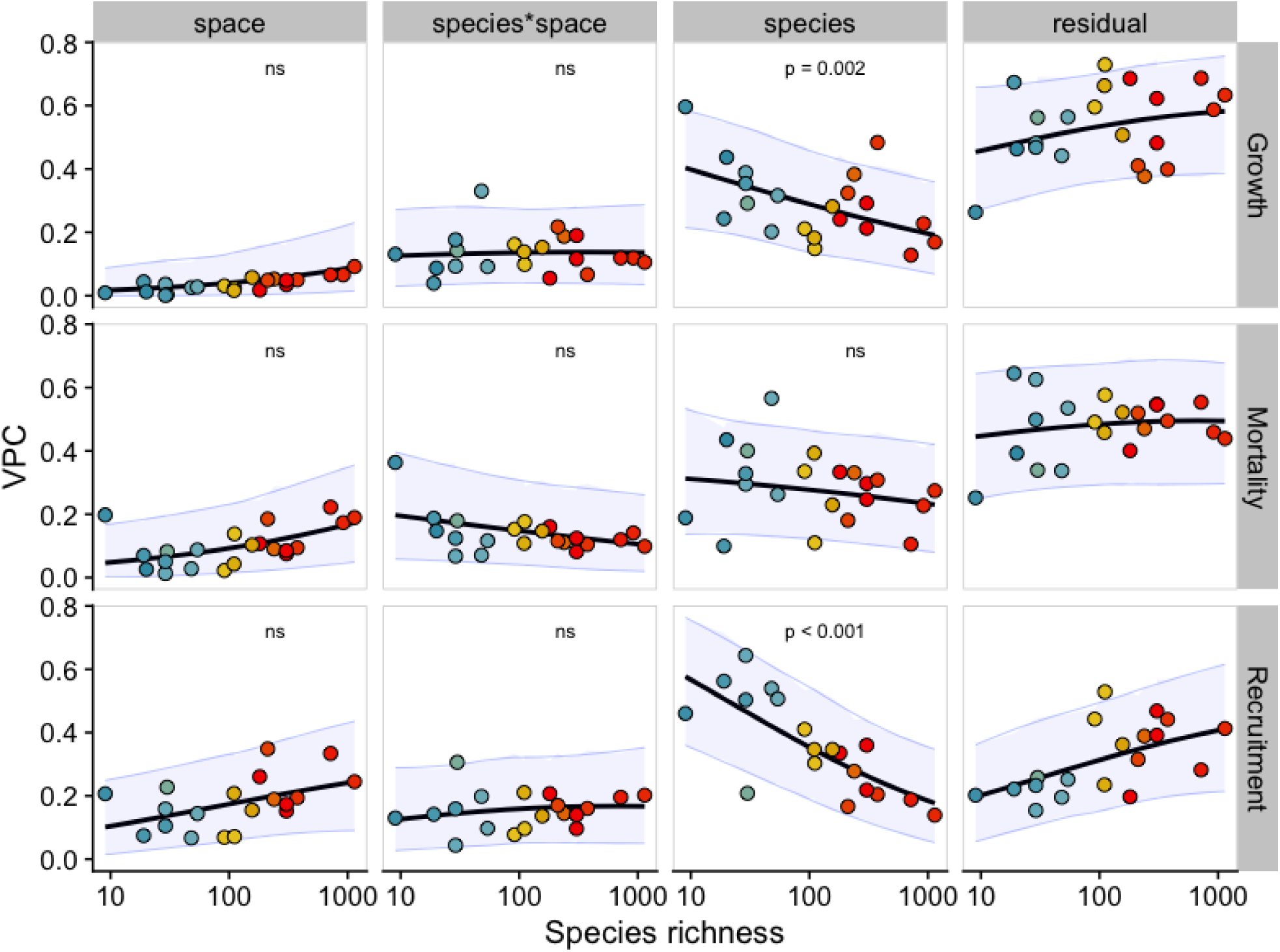
Variance partition coefficients (VPCs) for the organising principles (OPs) *species*, *space*, *species x space* and *residual* against species richness of 21 forest plots. OPs were estimated with the reduced model (eq. 1) without temporal OPs. Black lines are fitted relationships obtained from Dirichlet regressions of VPCs against species richness; shaded blue areas are the 95% prediction intervals. p-values are shown only for the significant values after Bonferroni correction (alpha=0.016). *Residual* VPCs are reference categories and thus were not tested for significance. Each forest plot (dots) is colored by absolute latitude as in Fig. 1a. Species richness on the x-axis is at the logarithmic scale with base 10.

## Discussion

Innumerable mechanisms operate and interact in forests and leave fingerprints of their integrated effects in tree vital rates, i.e., growth, survival, and recruitment, which together drive forest dynamics. Here, we used a conceptual and statistical framework to identify organising principles (OPs, Table 1) and quantify the associated variability among vital rates for more than 2.9 million trees of approx. 6,500 species in 21 forests across the globe. This, in turn, allows a first assessment of the relative importance of the groups of mechanisms underlying each OP, offering a first step in narrowing down which of the mechanisms are critical for structuring global forests. In the following sections, we summarise our most striking findings, discuss some potentially important mechanisms, and provide recommendations for an agenda to study tree vital rates.

### Species is a major source of variability in tree vital rates

We found that *species* was the most important OP for all tree vital rates, explaining on average between 29 and 36% of the demographic variance across all forest sites (Fig. 1). *Species* in interaction with *space* added another 13-15% variance explained, meaning that a total of 42-51% of demographic variation can be partitioned towards species differences and species-specific responses to spatial heterogeneity (Table 1). In contrast, *space* and *time* explained relatively little variability in vital rates (Fig. 1, 2). Our results, therefore, suggest that - at least at the temporal and spatial scales covered by our datasets - spatio-temporally varying factors alone contribute less to demographic variance than evolutionary history and adaptations to the environment. Grouping individuals into species thus creates a globally important cluster of demographic variation that appears consistently most important across a wide range of forests.

Our results on the importance of species support numerous ongoing research agendas. Efforts to include more realistic representation of species strategies in global vegetation models appear to be a promising route (Fisher *et al*., 2018; Anderegg *et al*., 2022), regardless of whether forest dynamics are studied in local tree neighbourhoods or larger spatial units (Fig. 3). We expect that accounting for species differences can explain up to ∼36% of demographic variation, while additionally accounting for small-scale species–environment associations (Messier et al., 2010; Lasky et al., 2014) might further improve this to almost half of the variation explained. More critically, however, our work shows that there are clear limits to the improvement more realistic representations of species can bring. Studies including species strategies typically rely on functional traits (Rubio & Swenson, 2022) or demographic trade-offs (Rüger *et al*., 2020; Russo *et al*., 2021), i.e., simplifications that explain only about half of the among-species variation (e.g., Visser *et al*., 2016). Nevertheless, the global importance of species in clustering demographic variance and its consistency across spatial scales indicates that endeavours seeking to better map species differences may have been undervalued compared to those focussing on spatial and temporal effects.

### Temporal variability acts mostly on recruitment and mortality and in interaction with space

In contrast to variability among species, temporal OPs played a minor role for variability in tree vital rates, as time interval alone was responsible for only 3-7% of total variability for plots with sufficient data. Although these data probably have the most comprehensive temporal coverage of large forest areas currently available, our findings might reflect the relatively short time frame (20 to 40 years), the low temporal resolution (approximately five years), and the fact that we could only analyse data from five tropical and subtropical forests. Nevertheless, variability between census intervals was detected in recruitment and to a lesser degree in mortality but was rather unimportant for growth (Fig. 2). A possible explanation is that growth rates fluctuate within shorter periods than our 5-year census interval can capture (Dobbertin, 2005), while recruitment and mortality may exhibit several bad or good years in a row (Schwartz *et al*., 2020).

Temporal effects were most important in interaction with space which, for instance, could indicate gap dynamics that jointly affect vital rates of most trees (Kohyama, 1993). This interpretation is consistent with the result that the *space x time* interaction OP was more important for mortality and recruitment than for growth - as mortality in gaps is known to be “spatially contagious” with falling trees killing multiple neighbours (Araujo *et al*., 2021), and the resulting gaps generally favour recruitment for many species (Brokaw, 1987). Additionally, some of the variability in the *space x time* OP could be the result of climatic events acting differently depending on local conditions, such as droughts that harm trees more in valleys than on ridges (Zuleta *et al*., 2017).

Our results on temporal OPs support a research agenda that should analyse the importance of climatic and/or temporal effects on vital rates in interaction with spatial effects. Moreover, we advocate for datasets with higher temporal resolution and longer time series, which would allow to capture larger but infrequent disturbances (Šamonil *et al*., 2013), thereby revealing more of the demographic importance of environmental fluctuations and temporal niches (Fung *et al*., 2020).

### Small spatial grain variability is important for tree vital rates

Spatial OPs were important for vital rate variability mostly in interaction with *species* for growth, and *time* for mortality and recruitment (Fig. 1 and 2), indicating the importance of spatial niches and patch dynamics (see previous section). Alone, *space* was the least important OP and only created considerable variability in models without *time* (Fig. 1). However, it may be possible that some spatial variability could still be present in the residual variance, since we used a simple, discrete spatial structure without accounting for spatial autocorrelation or more sophisticated spatial analysis.

Spatially acting mechanisms were best detected by dividing the plots into quadrats of 5x5 m (Fig. 3), indicating that trees interact and respond to local conditions at scales of a few metres through local mechanisms such as gap dynamics, competition, crown damage, and micro-topography (Schwartz *et al*., 2020). Further decreasing the spatial grain would then move below the scale of tree crowns, and begin to merely assign quadrats to single trees, here reflected by residual variance. With increasing grain size, less variability is explained by spatial mechanisms (Cáceres *et al*., 2012). Consequently, vital rates become less predictable at larger spatial grain. Nevertheless, even at the largest quadrat size of 100x100 m, spatial OPs still explained a reasonable part of the variability, with the consequence that tree species seem to distinctly respond also to environmental heterogeneity over larger areas (de Knegt *et al*., 2010), probably due to topography, water resources and soil nutrients (Russo *et al*., 2005, 2008; Zuleta *et al*., 2020).

### Large proportion of unexplained variability in tree vital rates

Residual variance was consistently the dominant component of the vital rate VPCs across sites and in the temporal and spatial analyses. In our variance partitioning analyses, residual variance represents the variance in the response that cannot be attributed to any of the grouping factors (here, the OPs). On one hand, this result encourages more detailed models that might include covariates that ‘explain’ differences among individual trees. For instance, both growth and mortality are known to differ across ontogeny, and thus tree size (e.g. dbh) should be able to explain some of the residual variance (Hülsmann *et al*., 2018). Moreover, functional traits at the individual level (Su *et al*., 2020) and structures that explicitly deal with spatial (Wiegand *et al*., 2017) and temporal autocorrelation may explain additional differences in individual vital rates. On the other hand, there are intrinsic limits to what can be explained by even the most detailed models, as the residual variance also includes inherent noise. This noise is the result of misattribution of species, mapping error or measurement error (Detto *et al*., 2019) and chaotic behaviour known to exist in many biological systems (Benincà *et al*., 2015).

### Globally, variability among species declines with species richness

Across plots, increasing species richness was associated with decreasing relative importance of the *species* OP in growth and recruitment (Fig. 4). This trend was robust to one of the most probable sources of bias, i.e., differences in species rarity between forest plots. Although species richness can strongly correlate with other environmental drivers (e.g., latitude, rainfall, biogeography), we consider that the decreasing relative importance of the *species* OP with species richness reflects a true macroecological pattern that could be further explored. Moreover, the decrease in the *species* VPC was determined by a decrease in the respective variance estimates, and not by an increase of variances related to the other OPs (Fig. S5.4). Similarly, Condit et al. (2006) found across ten tropical forests (seven in common with this study) that the range of species-specific mortality and growth rates decreased with higher species richness.

These results underpin that - in contrast to expectations of niche theory - the most diverse forests feature the lowest interspecific variation in vital rates. Following the rationale of niche theory, diverse forests should have more demographic niches than low-diversity forests, as more niches allow more species to have equivalent fitness thus favouring species coexistence (Chesson, 2000). The lack of evidence for wider demographic ranges in species-rich forests (this study, Condit *et al*., 2006; Clark, 2010) suggests that demographic niches play a minor role for large-scale diversity patterns, hinting towards more neutral dynamics (Hubbell, 2006). However, coexistence is inherently high dimensional, and comparing mean species values across low dimensions (a few vital rates) only partly represents the full niche space (Clark, 2010).

## Conclusions

As the mechanisms that influence vital rates can be grouped by the dimensions at which they operate and interact, patterns of how variance is partitioned along key dimensions can reveal how important various biotic and abiotic mechanisms are in influencing tree demography and hence forest dynamics. Here, we have shown that variance partitioning of vital rates among key ecological dimensions, i.e., species, space, and time, has the potential to provide a first step in identifying the structuring processes of global forest dynamics. We found that species differences were a major source of variability in tree vital rates, while temporal variability acted mostly on recruitment and in interaction with spatial variability. Small grain sizes captured most of the spatial variability, but there were still larger proportions of unexplained variability in vital rates, probably due to individual variation. Most intriguing, we found that, globally, variability among species declined with species richness. In summary, species in highly diverse forests present redundant vital rates that do not add to the diversity of demographic types, highlighting the challenges of studying and predicting changes in hyper diverse systems.

The proposed framework highlights the most promising avenues for future research both in terms of understanding the relative contributions of groups of mechanisms to forest demography and diversity, and for predicting forest ecosystems. We hope future studies may benefit from using this approach as a conceptual and modelling approach to narrow down which of the mechanisms are critical for structuring global forests.

## Acknowledgments

The authors thank the many people involved in establishing and maintaining all the plots utilised in these analyses. A detailed list of each forest plot funding sources, fieldwork permissions, acknowledgements, and references is available in Appendix S1. The development of this project benefited from the ForestGEO workshop 2019 in Singapore, organised by the Smithsonian Institution. Contributions by MSL were supported by travel grants from ForestGEO network (2020), University of Regensburg (2020), University of Bayreuth (2022), and PROEX-CAPES (Coordenação de Aperfeiçoamento de Pessoal de Nível Superior - Brazil, 2022). MSL received a 6-months Smithsonian Predoctoral Fellowship at the Smithsonian Ecological Research Center (2020-2021). LH received funding from the Bavarian Ministry of Science and the Arts within the Bavarian Climate Research Network (bayklif).

## Author contributions

MSL, SM, MV, LH, PIP, AOA conceived the ideas and conceptualization of the study.

MSL, SM, MV, LH conceived and designed the analyses.

MSL, LH curated the data. MSL performed the analyses.

MSL, SM, MV, LH wrote the initial draft.

MSL, SM, MV, LH, PIP, SD, HD wrote reviewed versions of the draft.

SA, KAA, NA, NAB, WYB, NC, CHCY, YYC, GC, KC, AD, SE, CENE, GG, IAUNG, CVSG, RH, WHH, AI, DJJ, DK, KK, YTL, JAL, JRM, YM, WJMS, MBM, MN, AN, GP, ReP, RoP, RPP, PS, IFS, ST, DT, JT, MU, AW, JZ, DZ contributed data and provided site-specific information

All authors contributed to the final version of the manuscript.

## Data archiving statement

The forest data that support the findings of this study are available from the ForestGEO network. For some of the sites, the data is publicly available at https://forestgeo.si.edu/explore-data. Restrictions apply, however, to the availability of the data from other sites, which were used under license for the current study, and so are not publicly available. Data are however available from the authors upon reasonable request and with permission of the principal investigators of the ForestGEO sites. We provide an example of data preparation and analysis workflow from a forest plot with public available data and the code for all results and analyses on Zenodo repository (Leite, 2022).

## Ethics statement

Licenses and permissions for the fieldwork and monitoring of the ForestGEO Forest Dynamic Plots are provided separately for each forest plot in the section Forest plots acknowledgments and references in Appendix S1.

