## Supplementary material for "Major axes of variation in tree demography across global forests"

#### Appendix S1 - Forest census data

##### Forest plots data preparation

In every forest plot dataset, each observation is an individual tree. For trees with more than one stem, the individual is considered alive if at least one of the stems is alive and dead if all the stems are dead. The diameter at breast height (DBH) for trees with multiple stems was calculated based on the sum of the basal area of the stems, considering the diameter of a circle.

We excluded ferns and palms from all analyses due to their non-standard growth form, and lack of secondary growth. We also excluded trees without information on *x* and *y* plot coordinates, species name, status (alive, dead or recruit), and date of measurement. Although we excluded individuals with unknown or unidentified species name, we kept morphospecies classification when existing in the data. For growth analysis, we excluded individuals with different heights of measurements of DBH in consecutive censuses and trees with growth rates more than four standard deviations from the mean, as they are likely measurement errors (Rüger *et al.* 2011; Condit *et al.* 2017).

Environmental, climatic and vegetational information of each ForestGEO plot are in Table S1.1 and the summary information of the data used in the analysis in Table S1.2.

**TABLE S1.1:** ForestGEO plots information. Data on environmental and climatic variables are from Anderson-Teixeira *et al.* (2015). Abbreviations and units in columns: Köppen Climate classification zone*; MAT mean annual temperature in °C; MAP mean annual precipitation in mm/year; PET annual potential evapotranspiration in mm/day; Dominant soil Classification**; Dominant vegetation type*** Natural disturbance regime****.

| Forest Plot | Latitudinal zone | Country | Latitude | Longitude | Elevation m (min-max) | Köppen | MAT | MAP | PET | Dominant soil | Dominant vegetation | Natural disturbances |
|---|---|---|---|---|---|---|---|---|---|---|---|---|
| Amacayacu | Tropical | Colombia | -3.81 | -70.27 | 89-111 | Af | 25.8 | 3216 | 1010 | Ult | BE | FI; W; In |
| Barro Colorado Island | Tropical | Panama | 9.15 | -79.85 | 120-160 | Am | 27.1 | 2551 | 1311 | Ox | BdD; BE | D; W |
| Fushan | Subtropical | Taiwan | 24.76 | 121.56 | 600-733 | Cfa | 18.2 | 4271 | 1085 | Ult; In | BE | H |
| Ilha do Cardoso | Subtropical | Brazil | -25.096 | -47.9573 | 3-8 | Cfa | 22.4 | 2100 | - | S | BE | - |
| Ituri - Edoro | Tropical | Democratic Republic of Congo | 1.44 | 28.583 | 700-850 | Af | 24.3 | 1682 | 1168 | Ox | BE | W; A |
| Ituri - Lenda | Tropical | Democratic Republic of Congo | 1.44 | 28.583 | 700-850 | Af | 24.3 | 1682 | 1168 | Ox | BE | W; A |
| Korup | Tropical | Cameroon | 5.07 | 8.85 | 150-240 | Am | 26.6 | 5272 | 1050 | Ult; Ox | BE | W |
| Lambir | Tropical | Malaysia | 4.19 | 114.02 | 104-244 | Af | 26.6 | 2664 | 1114 | Ult | BE | L; D |
| Lilly Dickey Woods | Temperate | USA | 39.24 | -86.22 | 230-303 | Cfa | 11.6 | 1203 | 981 | In; Ult; Alf | BcD | W; D; Ic |
| La Planada | Tropical | Colombia | 1.16 | -77.99 | 1796-1840 | Cfb | 19 | 4087 | - | An | BE | W |
| Luquillo | Tropical | Puerto Rico, USA | 18.33 | -65.82 | 333-428 | Af | 22.8 | 3548 | 1219 | Ox; Ult | BE | H; L |
| Mo Singto | Tropical | Thailand | 14.43 | 101.35 | 725-815 | Aw | 23.5 | 2100 | 1300 | NA | BE; BdD | W |
| Pasoh | Tropical | Malaysia | 2.98 | 102.31 | 70-90 | Af | 27.9 | 1788 | 1120 | Ult | BE | W |
| Smithsonian Conservation Biology Institute | Temperate | USA | 38.89 | -78.15 | 273-338 | Cfa | 12.9 | 1001 | 1003 | Alf | BcD | W, Ic |
| Smithsonian Environmental Research Center | Temperate | USA | 38.89 | -76.56 | 6-10 | Cfa | 13.2 | 1068 | 1111 | Ult; In; En | BcD | H; W |

| Forest Plot | Latitudinal zone | Country | Latitude | Longitude | Elevation m (min-max) | Köppen | MAT | MAP | PET | Dominant soil | Dominant vegetation | Natural disturbances |
|---|---|---|---|---|---|---|---|---|---|---|---|---|
| Sinharaja | Tropical | Sri Lanka | 6.40 | 80.40 | 424-575 | Af | 22.5 | 5016 | 1384 | Ult | BE | W |
| University of California Santa Cruz | Temperate | USA | 37.01 | -122.08 | 314-332 | Csb | 14.8 | 778 | - | Mo | BcD | W, Ic |
| Wabikon | Temperate | USA | 45.55 | -88.79 | 488-514 | Dfb | 4.2 | 805 | - | Alf | BdC | W |
| Wind River | Temperate | USA | 45.82 | -121.96 | 352-385 | Csb | 9.2 | 2495 | 770 | An | NE | Fi; W; In |
| Wytham Woods | Temperate | United Kingdom | 51.77 | -1.34 | 104-163 | Cfb | 10 | 717 | 637 | E | BcD | - |
| Zofin | Temperate | Czech Republic | 48.66 | 14.71 | 735-825 | Cfb | 6.2 | 866 | - | S; In; Hi | BdC; NE | W; In |

*Af: Tropical with significant precipitation year-round; Am: Tropical monsoon; Aw: Tropical wet and dry; Csb subtropical/mid-latitude climate with dry summers (a.k.a.: Warm-summer Mediterranean); Cfa: Humid subtropical/mid-latitude climate with significant precipitation year-round; Cfb: Oceanic with significant precipitation year-round; Dfb: Humid Continental with significant precipitation year-round.
** Alf, Alfisols; An, Andisols; E, Entisoils; Ge, Gelisols; Hi, Histosols; In, Inceptisols; Ox, Oxisols; Ult, Ultisols; S, Spodosols; Ve, Vertisols.
*** BE, broadleaf evergreen; BdD, broadleaf drought deciduous; BcD, broadleaf cold deciduous; NE, needleleaf evergreen.
****A, animal activity (destructive); D, Drought; E, Erosion; Fi, Fire; Fl, flood; H, hurricane/typhoon; Ic, Ice storms; Insect outbreaks; L, landslides; PT, permafrost thaw; W, wind storms (local).

**TABLE S1.2.** Summary information of the data used in the analysis. Number of species may include morphospecies.

| ID | Forest Plot | Original/ Trimmed plot size (ha) | First census (year) | Last census (year) | Number of census intervals | Total Number of species | | | Total number of observations | | |
|---|---|---|---|---|---|---|---|---|---|---|---|
| | | | | | | Growth | Mortality | Recruitment | Growth | Mortality | Recruitment |
| ama | Amacayacu | 25/25 | 2007 | 2017 | 1[+] | 1156 | 1269 | - | 76580 | 105357 | - |
| bci | Barro Colorado Island | 50/50 | 1981 | 2016 | 7* | 305 | 313 | 316 | 1310125 | 1558414 | 1620133 |
| fus | Fushan | 25/25 | 2003 | 2019 | 3 | 105 | 107 | 105 | 267550 | 325882 | 341664 |
| idc | Ilha do Cardoso | 10.24/9 | 2009 | 2019 | 1 | 116 | 117 | 130 | 17529 | 19081 | 25306 |
| edo | Ituri - Edoro | 20/20 | 1994 | 2007 | 2 | 388 | 412 | 417 | 280412 | 313806 | 305282 |
| len | Ituri - Lenda | 20/20 | 1994 | 2007 | 2 | 382 | 396 | 399 | 234991 | 264914 | 259444 |
| kor | Korup | 50/50 | 1997 | 2010 | 1 | 449 | 468 | 461 | 280703 | 327121 | 321582 |
| lam | Lambir | 52/50 | 1991 | 2009 | 3 | 1362 | 1402 | 1376 | 915971 | 1073643 | 1065051 |
| ldw | Lilly Dickey Woods | 25/25 | 2012 | 2017 | 1 | 33 | 36 | 33 | 20596 | 26496 | 23059 |
| lpl | La Planada | 25/25 | 1997 | 2003 | 1 | 203 | 205 | 225 | 71713 | 89251 | 89359 |
| luq | Luquillo | 16/15 | 2001 | 2016 | 3 | 134 | 146 | 145 | 106355 | 166558 | 156824 |
| mos | Mo Singto | 30.5/30 | 2003 | 2017 | 2 | 266 | 272 | 275 | 227085 | 273510 | 394809 |
| pas | Pasoh | 50/50 | 1986 | 2011 | 5* | 880 | 891 | 891 | 1357235 | 1555377 | 1588440 |
| scbi | Smithsonian Conservation Biology Institute | 25.6/24 | 2008 | 2013 | 1 | 57 | 66 | 60 | 24265 | 29022 | 33812 |
| serc | Smithsonian Environmental Research Center | 16/16 | 2008 | 2014 | 1 | 65 | 71 | 70 | 19834 | 23200 | 24156 |
| sin | Sinharaja | 25/25 | 1993 | 2008 | 2 | 231 | 234 | 231 | 355214 | 399749 | 381212 |
| ucsc | University of California | 16/6 | 2006 | 2020 | 2 | 30 | 31 | 30 | 12873 | 15753 | 14634 |

| ID | Forest Plot | Original/ Trimmed plot size (ha) | First census (year) | Last census (year) | Number of census intervals | Total Number of species | | | Total number of observations | | |
|---|---|---|---|---|---|---|---|---|---|---|---|
| | | | | | | Growth | Mortality | Recruitment | Growth | Mortality | Recruitment |
| | Santa Cruz | | | | | | | | | | |
| wab | Wabikon | 25.6/24 | 2008 | 2018 | 2 | 33 | 36 | 34 | 77167 | 92258 | 84133 |
| wfdp | Wind River | 27.2/24 | 2010 | 2016 | 1 | 24 | 26 | 25 | 22979 | 25354 | 24420 |
| wyw | Wytham Woods | 18/18 | 2008 | 2021 | 3[+] | 25 | 25 | - | 54198 | 57925 | - |
| zof | Zofin | 25/25 | 2012 | 2017 | 1 | 11 | 11 | 12 | 57445 | 58344 | 72764 |

*for growth rates, Barro Colorado Island had 6 and Pasoh 4 census intervals due to problems with DBH measurements in the first census.
[+] Recruitment rates for Amacayacu and Wytham Woods could not be analysed. Wytham Woods was not analysed with temporal data.

**References**

Anderson-Teixeira, K.J., Davies, S.J., Bennett, A.C., Gonzalez-Akre, E.B., Muller-Landau, H.C., Joseph Wright, S., Abu Salim, K., Almeyda Zambrano, A.M., Alonso, A., Baltzer, J.L., Basset, Y., Bourg, N.A., Broadbent, E.N., Brockelman, W.Y., Bunyavejchewin, S., Burslem, D.F.R.P., Butt, N., Cao, M., Cardenas, D., Chuyong, G.B., Clay, K., Cordell, S., Dattaraja, H.S., Deng, X., Detto, M., Du, X., Duque, A., Erikson, D.L., Ewango, C.E.N., Fischer, G.A., Fletcher, C., Foster, R.B., Giardina, C.P., Gilbert, G.S., Gunatilleke, N., Gunatilleke, S., Hao, Z., Hargrove, W.W., Hart, T.B., Hau, B.C.H., He, F., Hoffman, F.M., Howe, R.W., Hubbell, S.P., Inman-Narahari, F.M., Jansen, P.A., Jiang, M., Johnson, D.J., Kanzaki, M., Kassim, A.R., Kenfack, D., Kibet, S., Kinnaird, M.F., Korte, L., Kral, K., Kumar, J., Larson, A.J., Li, Y., Li, X., Liu, S., Lum, S.K.Y., Lutz, J.A., Ma, K., Maddalena, D.M., Makana, J.-R., Malhi, Y., Marthews, T., Mat Serudin, R., McMahon, S.M., McShea, W.J., Memiaghe, H.R., Mi, X., Mizuno, T., Morecroft, M., Myers, J.A., Novotny, V., de Oliveira, A.A., Ong, P.S., Orwig, D.A., Ostertag, R., den Ouden, J., Parker, G.G., Phillips, R.P., Sack, L., Sainge, M.N., Sang, W., Sri-ngernyuang, K., Sukumar, R., Sun, I.-F., Sungpalee, W., Suresh, H.S., Tan, S., Thomas, S.C., Thomas, D.W., Thompson, J., Turner, B.L., Uriarte, M., Valencia, R., Vallejo, M.I., Vicentini, A., Vrška, T., Wang, X., Wang, X., Weiblen, G., Wolf, A., Xu, H., Yap, S. & Zimmerman, J. (2015) CTFS-ForestGEO: a worldwide network monitoring forests in an era of global change. *Global Change Biology*, **21**, 528–549.

Condit, R., Pérez, R., Lao, S., Aguilar, S. & Hubbell, S.P. (2017) Demographic trends and climate over 35 years in the Barro Colorado 50 ha plot. *Forest Ecosystems*, **4**, 17.

Rüger, N., Berger, U., Hubbell, S.P., Vieilledent, G. & Condit, R. (2011) Growth Strategies of Tropical Tree Species: Disentangling Light and Size Effects. *PLoS ONE*, **6**, e25330.

#### Forest plots acknowledgments and references

##### Amacayacu

The 25-ha Long-Term Ecological Research Project of Amacayacu is a collaborative project of the Instituto Amazónico de Investigaciones Científicas Sinchi and the Universidad Nacional de Colombia Sede Medellín, in partnership with the Unidad de Manejo Especial de Parques Naturales Nacionales and the Forest Global Earth Observatory of the Smithsonian Tropical Research Institute (ForestGEO). The Amacayacu Forest Dynamics Plot is part of ForestGEO, a global network of large-scale demographic tree plots. We acknowledge the Director and staff of the Amacayacu National Park for supporting and maintaining the project in this National Park.

##### Barro Colorado Island

The BCI forest dynamics research project was made possible by National Science Foundation grants to Stephen P. Hubbell: DEB-0640386, DEB-0425651, DEB-0346488, DEB-0129874, DEB-

00753102, DEB-9909347, DEB-9615226, DEB-9615226, DEB-9405933, DEB-9221033, DEB-9100058, DEB-8906869, DEB-8605042, DEB-8206992, DEB-7922197, support from the Forest Global Earth Observatory, the Smithsonian Tropical Research Institute, the John D. and Catherine T. MacArthur Foundation, the Mellon Foundation, the Small World Institute Fund, and numerous private individuals, and through the hard work of over 100 people from 10 countries over the past three decades. The plot project is part of the Forest Global Earth Observatory (ForestGEO), a global network of large-scale demographic tree plots.

Plot references:

Condit R., Perez, R., Aguilar, S., Lao, S., Foster, R., Hubbell, S.P. 2019. Complete data from the Barro Colorado 50-ha plot: 423617 trees, 35 years, 2019 version.

Condit, R. 1998. Tropical Forest Census Plots. Springer-Verlag and R. G. Landes Company, Berlin, Germany, and Georgetown, Texas.

Hubbell, S.P., R.B. Foster, S.T. O'Brien, K.E. Harms, R. Condit, B. Wechsler, S.J. Wright, and S. Loo de Lao. 1999. Light gap disturbances, recruitment limitation, and tree diversity in a neotropical forest. Science 283: 554-557.

Condit R., Perez, R., Aguilar, S., Lao, S., Foster, R., Hubbell, S.P. 2019. BCI 50-ha plot taxonomy, 2019 version.

**Fushan**

The Fushan Forest Dynamics plot (FDP) is supported by the Taiwan Forestry Bureau, the Taiwan Forestry Research Institute and the Ministry of Science and Technology of Taiwan. We would like to express our gratitude to all field technicians and students who helped with the implementation and recensus of the Fushan FDP. We also thank the Fushan Research Center staff for providing logistic support.

**Ilha do Cardoso**

The 10.24ha Ilha do Cardoso Forest Dynamics Plot was established and has been supported by São Paulo Research Foundation (FAPESP grants #1999/09635-0 and #2017/11979-9), University of São Paulo (USP), Forest Global Earth Observatory and  the Smithsonian Tropical Research Institute. The Principal Investigator (AAdeO) received fellowship grant from Brazil National Council for Scientific and Technological Development (CNPq #313829/2021-7) and would like to express his gratitude to many field workers, research technicians, and staff who helped during decades, working at field and USP, to make this data available.

Plot references:

Oliveira, A.A. de, Vicentini, A., Chave, J., Castanho, C. de T., Davies, S.J., Martini, A.M.Z., Lima, R.A.F., Ribeiro, R.R., Iribar, A., and Souza, V.C. (2014). Habitat specialization and phylogenetic structure of tree species in a coastal Brazilian white-sand forest. Journal of Plant Ecology 7, 134–144. 10.1093/jpe/rtt073.

Lima, R.A.F. de, Oliveira, A.A. de, Martini, A.M.Z., Sampaio, D., Souza, V.C., and Rodrigues, R.R. (2011). Structure, diversity, and spatial patterns in a permanent plot of a high Restinga forest in Southeastern Brazil. Acta Bot. Bras. 25, 633–645. 10.1590/S0102-33062011000300017.

**Ituri - Edoro and Lenda**

The Ituri 40-ha plot program is a collaborative project between the Centre de Formation et de Recherche en Conservation Forestière, the Wildlife Conservation Society – DRC through his conservation project in the Okapi forest Reserve, in partnership with the Forest Global Earth Observatory (ForestGEO). The Ituri plots are financially supported by the Wildlife Conservation Society, the Frank Levinson Family Foundation, and ForestGEO. The Institut Congolais pour la Conservation de la Nature graciously provided the research permit.

Plot references:

Makana J-R., Hart T.B., Liengola I., Ewango C.N.E., Hart J.A. & Condit R. (2004). Ituri Forest Dynamics Plots, Democratic Republic of Congo. In Losos E.C. and Leigh, E.G. Jr. eds. Forest Diversity and Dynamism: Findings from a Large-Scale Plot Network. . 506-516. University of Chicago Press, Chicago.

**Korup**

The 50-ha is a collaborative project of the University of Buea, Cameroon, and the World Wide Fund for Nature, Cameroon Program in partnership with the Forest Global Earth Observatory of the Smithsonian Tropical Research Institute (ForestGEO). Funding for the first census was provided by the International Cooperative Biodiversity Group (a consortium of the NIH, the NSF, and the USDA), with supplemental funding by the Central Africa Regional Program for the Environment (a program of USAID). Funding for the second census was provided by the Frank Levinson Family Foundation. Permission to conduct the field program in Cameroon is provided by the Ministry of Environment and Forests and the Ministry of Scientific Research and Innovation.

Plot references:

Chuyong G.B., Condit, R., Kenfack, D., Losos, E., Sainge, M., Songwe, N.C., and Thomas, D.W. (2004). Korup Forest Dynamics Plot, Cameroon. In Losos E.C. and Leigh, E.G. Jr. eds. Forest Diversity and Dynamism: Findings from a Large-Scale Plot Network. . 506-516. University of Chicago Press, Chicago.

Thomas, D.W., Kenfack, D., Chuyong, G.B., Sainge N.M., Losos E.C., Condit R.S., Songwe N.C. (2003). Tree Species of Southwestern Cameroon: Tree distribution maps, diameter tables and species documentation of the 50-ha Korup Forest Dynamics Plot. Center for Tropical Forest Science, Washington, D.C.

Kenfack D., Thomas D.W., Chuyong G.B., & Condit R. (2007). Rarity and abundance in a diverse African forest: Biodiversity Conservation 16: 2045 – 2074.

**Lambir**

The 52-ha Long-Term Ecological Research Project is a collaborative project of the Forest Department of Sarawak, Malaysia, the Forest Global Earth Observatory (ForestGEO) , the Arnold Arboretum of Harvard University, USA (under NSF awards DEB-9107247 and DEB-9629601), and Osaka City, Ehime & Kyoto Universities, Japan (under MEXT/JSPS KAKENHI grants 09NP0901, 22H02388, and JST/JICA-SATREPS PUBS). The Lambir Forest Dynamics Plot is part of ForestGEO, a global network of large-scale demographic tree plots. We acknowledge the Sarawak Forest Department for supporting and maintaining the project in Lambir Hills National Park.

Plot references:

Lee, H.S., P.S. Ashton, T. Yamakura, S. Tan, S.J. Davies, A. Itoh, T. Ohkubo & J.V. LaFrankie. (2002). The 52-Hectare Forest Dynamics Plot at Lambir Hills, Sarawak, Malaysia: Tree Distribution Maps, Diameter Tables and Species Documentation.Sarawak Forest Department. 621 pp. Lee Ming Press, Kuching, Sarawak, Malaysia.

Lee, H., Davies, S. J., LaFrankie, J. V., Tan, S., Itoh, A., Yamakura, T., and Ashton, P. S. 2002. Floristic and structural diversity of 52 hectares of mixed dipterocarp forest in Lambir Hills National Park, Sarawak, Malaysia. Journal of Tropical Forest Science, 14: 379-400.

**La Planada**

The 25-ha is a collaborative project between the Instituto de Investigación de Recursos Biológicos Alexander von Humboldt and the Forest Global Earth Observatory (ForestGEO) of the Smithsonian Tropical Research Institute. We thank Natalia Norden Medina for making the data available. And La Planada forest plot especially thanks to Martha Isabel Vallejo and Cristian Samper, who made this project possible. For more information on La Planada, visit:
http://i2d.humboldt.org.co/ceiba/resource.do?r=planada_parcelapermanente_censo1

**Lilly Dickey Woods**

The 25-ha Indiana University Forest Dynamics Plot is a collaborative project of Indiana University and the Center for Tropical Forest Science of the Smithsonian Tropical Research Institute. Funding for the installation and maintenance of the IUFDP came from multiple sources including Indiana Academy of Science, Indiana University Research and Teaching Preserve, U.S. Department of Energy, the USDA National Institute for Food and Agriculture McIntire Stennis project 1018790, and ForestGEO. The IUFDP is part of ForestGEO, a global network of large-scale demographic tree plots.

Plot references:

Johnson, D.J., Clay, K. & Phillips, R.P. Mycorrhizal associations and the spatial structure of an old-growth forest community. Oecologia 186, 195–204 (2018). https://doi.org/10.1007/s00442-017-3987-0

**Luquillo**

This research was supported by grants BSR-8811902, DEB 9411973, DEB 0080538, DEB 0218039, DEB 0620910, DEB 0963447 and DEB-129764 from NSF to the Department of Environmental Science, University of Puerto Rico, and to the International Institute of Tropical Forestry, USDA Forest Service, as part of the Luquillo Long-Term Ecological Research Program. Thanks to the Andrew Mellon Foundation for funding. The U.S. Forest Service (Dept. of Agriculture) the University of Puerto Rico gave additional support. Thanks to all of the volunteers and staff members that have censused the plot. The LFDP is part of the Smithsonian Institution Forest Global Earth Observatory, a worldwide network of large, long-term forest dynamics plots.

Plot references:

Zimmerman, Jess K., Liza S. Comita, Jill Thompson, María Uriarte, and Nicholas Brokaw. 2010. Patch dynamics and community metastability of a subtropical forest: compound effects of natural disturbance and human land use. Landscape Ecology 25: 1099-1111.

Thompson, J., Brokaw, N., Zimmerman, J.K., Waide, R.B., Everham, III, E.M., Lodge, D.J., Taylor, C.M., García-Montiel, D. and Fluet, M. 2002. Land use history, environment, and tree composition in a tropical forest. Ecological Applications 12, 1344-1363.

**Mo Singto**

The 30.5-ha Mo Singto Forest Dynamics Plot is supported by Mahidol University, National Center for Genetic Engineering and Biotechnology, National Science and Technology Development Agency, and Department of National Parks, Wildlife and Plant Conservation. Many thanks to plot Principal Investigators Anuttara Nathalang and Warren Y. Brockelman, and to countless field workers, research and data technicians, and staff.

**Pasoh**

Data from the Pasoh Research Forest was provided by the Forest Research Institute Malaysia - Forest Global Earth Observatory, Smithsonian Tropical Research Institute collaborative research project. Negeri Sembilan Forestry Department is the custodian of Pasoh Research Forest and we acknowledge the department for preserving the research forest.

Plot references:

Manokaran N & LaFrankie JV. 1990. Stand structure of Pasoh Forest Reserve, A lowland rainforest in Peninsular Malaysia. Journal of Tropical Forest Science 3: 14-24.

Kochumen KM, LaFrankie JV, Manokaran N. 1990. Floristic composition of Pasoh Forest Reserve, a lowland forest in Peninsular Malaysia. Journal of Tropical Forest Science 3: 1-13.

Manokaran N, Abd Rahman K, Azman H, Quah ES, Chong PF. 1992. Short-term population dynamics of dipterocarp trees in a lowland rainforest in Peninsular Malaysia. Journal of Tropical Forest Science 5: 97-112.

**Smithsonian Conservation Biology Institute - SCBI**

Funding for the Smithsonian Conservation Biology Institute (SCBI) Large Forest Dynamics Plot (LFDP) was provided by the Smithsonian Institution, the National Zoological Park, and the HSBC Climate Partnership. The SCBI LFDP is part of the Smithsonian Institution Forest Global Earth Observatory, a worldwide network of large, long-term forest dynamics plots.

Plot references:

Bourg, N.A., W.J. McShea, J.R. Thompson, J.C. McGarvey, and X. Shen. 2013. Initial census, woody seedling, seed rain, and stand structure data for the SCBI SIGEO Large Forest Dynamics Plot. Ecology 94(9): 2111-2112. http://dx.doi.org/10.1890/13-0010.1

**Smithsonian Environmental Research Center - SERC**

Data on SERC Dynamic Forest plot was provided by Geoffrey Parker on October 8, 2020. These data were gathered as part of forest ecology studies at the Smithsonian Environmental Research Center (SERC). SERC is a participant in the Smithsonian Institution Forest Global Earth Observatory (ForestGEO) network.

**Sinharaja**

The 25-ha Long-Term Ecological Research Project at Sinharaja World Heritage Site is a collaborative project of the Uva Wellassa University, University of Peradeniya, the Forest Global Earth Observatory (ForestGEO) of the Smithsonian Tropical Research Institute, with supplementary funding received from the John D. and Catherine T. Macarthur Foundation, the National Institute for Environmental Science, Japan, and the Helmholtz Centre for Environmental Research-UFZ, Germany, for past censuses. The PIs gratefully acknowledge the Forest Department, Uva Wellassa University, and the Post-Graduate Institute of Science at the University of Peradeniya, Sri Lanka for supporting this project, and the local field and lab staff who tirelessly contributed in the repeated censuses of this plot.

**University of California Santa Cruz - UCSC**

The UCSC Forest Ecology Research Plot was made possible by National Science Foundation grants to Gregory S. Gilbert (DEB-0515520, DEB-084259, and DEB-1655896), by the Pepper-Giberson Chair Fund, the Robert Headley Presidential Chair for Integral Ecology and Environmental Justice, the UCSC Campus Natural Reserve, the University of California, the ForestGEO global network, and by the hard work of hundreds of UCSC students. We acknowledge that the land on which this research was conducted is the unceded territory of the Awaswas-speaking Uypi Tribe. The Amah Mutsun Tribal Band, comprised of the descendants of indigenous people taken to missions Santa Cruz and San Juan Bautista during Spanish colonization of the Central Coast, is today working hard to restore traditional stewardship practices on these lands and heal from historical trauma.

Plot references:

Gilbert, G.S., E. Howard, B. Ayala-Orozco, M. Bonilla-Moheno, J. Cummings, S. Langridge, I.M. Parker, J. Pasari, D. Schweizer, S. Swope. 2010. Beyond the tropics: forest structure in a temperate forest mapped plot. Journal of Vegetation Science 21: 388-405.

**Wabikon**

The Wabikon Lake Forest Dynamics Plot, located in the Chequamegon-Nicolet National Forest of northern Wisconsin, is part of the Smithsonian Institution's ForestGEO network. Tree censuses at the site have been supported by The 1923 Fund, the Smithsonian Tropical Research Institute, and the Cofrin Center for Biodiversity at the University of Wisconsin-Green Bay. More than 50 scientists and student and assistants contributed to the first three plot censuses. We are particularly grateful for the important contributions by Gary Fewless, Steve Dhein, Kathryn Corio, Juniper Sundance, Cindy Burtley, Curt Rollman, Mike Stiefvater, Kim McKeefry, Lukas Magee, Jon Schubbe, and U.S. Forest Service collaborators Linda Parker and Steve Janke.

Plot reference:

Magee, L., A. Wolf, R. Howe, J. Schubbe, K. Hagenow, and B. Turner. 2021. Density dependence and habitat heterogeneity regulate seedling survival in a North American temperate forest. Forest Ecology and Management 480: 118722.

**Wind River**

The Wind River Forest Dynamics Plot is a collaborative project of Utah State University, the Utah Agricultural Experiment Station and the USDA Forest Service Pacific Northwest Research Station. Funding was provided by the Center for Tropical Forest Science of the Smithsonian Tropical Research Institute, Utah State University, and the Utah State Agricultural Experiment Station. We acknowledge the Gifford Pinchot National Forest and the Wind River Field Station for providing logistical support, and the students, volunteers and staff individually listed at http://wfdp.org for data collection.

Plot references:

Lutz, J. A, A. J. Larson, J. A. Freund, M. E. Swanson, and K. J. Bible. 2013. The ecological importance of large-diameter trees to forest structural heterogeneity. PLOS ONE 8(12): e82784.

Lutz, J. A., A. J. Larson, T. J. Furniss, J. A. Freund, M. E. Swanson, D. C. Donato, K. J. Bible, J. Chen, and J. F. Franklin. 2014. Spatially non-random tree mortality and ingrowth maintain equilibrium pattern in an old-growth *Pseudotsuga-Tsuga* forest. *Ecology* 95(8): 2047-2054.

**Wytham Woods**

The 18-ha Long-Term Forest Monitoring Plot is a collaborative project between the University of Oxford, the Centre for Ecology and Hydrology, and the Smithsonian Institution ForestGEO (HSBC

Climate Partnership). The Wytham Forest Monitoring Plot is part of ForestGEO, a global network of large-scale demographic tree plots. Censuses were funded with support from ForestGEO and Advanced Investigator award from European Research Council to YM (GEM-TRAIT).

Plot references:

Butt, N., Campbell, G., Malhi, Y., Morecroft, M., Fenn, K., Thomas, M. (2009) Initial results from establishment of a long-term broadleaf monitoring plot at Wytham Woods, Oxford, UK. University of Oxford Report.

**Zofin**

The Zofin Forest Dynamics Plot was established with the support of the Smithsonian Institution as a part of the Smithsonian Institution Forest Global Earth Observatory, a worldwide network of large, long-term forest dynamics plots. We acknowledge the Department of Forest Ecology of the Silva Tarouca Research Institute for supporting and maintaining the long-term monitoring of the Zofin Forest Dynamics Plot (under the Czech Science Foundation, grant No. 19-09427S).

Plot references:

Janík D., Král K., Adam D., Hort L., Šamonil P., Unar P., Vrška T., McMahon S.M., 2016. Tree spatial patterns of Fagus sylvatica expansion over 37 years. Forest Ecology and Management 375: 134–145.

Janík D., Vrška T., Hort L., Unar P., Král K., 2018. Where have all the tree diameters grown? Patterns in Fagus sylvatica L. diameter growth on their run to the upper canopy. Ecosphere 9 (12): 1-19.

Šamonil, P., Daněk, P., Lutz, J.A., Anderson-Teixeira, K.J., Jaroš, J., Phillips, J.D., Rousová, A., Adam, D., Larson, A.J., Kašpar, J., Janik, D., Vašíčková, I., Gonzalez-Akre, E. & Egli, M. (2022) Tree Mortality may Drive Landscape Formation: Comparative Study from Ten Temperate Forests. *Ecosystems*. https://doi.org/10.1007/s10021-022-00755-8.

### Appendix S2 - Robustness analysis for subsampling forest plots

#### Comparing forest plots of different sizes

Because forest plots differed in size (from 6 to 50 ha), we tested if differences in plot size could bias VPC estimates. For that, we compared VPC of the reduced model (without temporal OPs) from Lambir using the entire plot with the average VPCs from 10 subsets of 5 ha each. Lambir is a suitable plot for such a comparison because it has one of the largest species richness (> 1000) and it is a large plot size (50 ha). We randomly subsampled Lambir data 10 times to the size of 5 ha and ran reduced models for growth at the 5x5 m quadrat size to build distributions of the estimates and compared mean values with the estimates of the results for the 50 ha analysis.

We also evaluate if the approach of repeatedly fitting the vital rate models to smaller subsamples of the plot, which was necessary to run the models with temporal OPs, is robust. For this purpose, we ran the VPC analysis (eq.1) for the entire Fushan (25 ha) and Luquillo (15 ha) forest plots, at the 100x100 m quadrat size for each vital rate and compared VPC estimates against the distribution of VPC estimates obtained from 10 subsampled datasets of 5 ha at the same quadrat size.

For both robustness analyses, using subsamples of the entire plot only marginally changed the estimates and, thus, we conclude that (1) the results among forests with different plot sizes for models without temporal OPs can be compared, and (2) the average VPCs among subsampled datasets are reliable VPC estimates for models with temporal OPs.

#### Models without temporal OPs - Lambir forest plot

The 5 ha subsamples presented on average 86% of the original number of species (1033 of 1311) and 10% of the original number of trees (mean 32,007 of 313,544). Mean VPCs from models using the subsamples of 5 ha were very similar to VPC values calculated from the entire 50 ha forest data (Figure S2.1), being negligible for *space* and *species x space* OPs and around 0.02 for *species* and *residual* OPs.

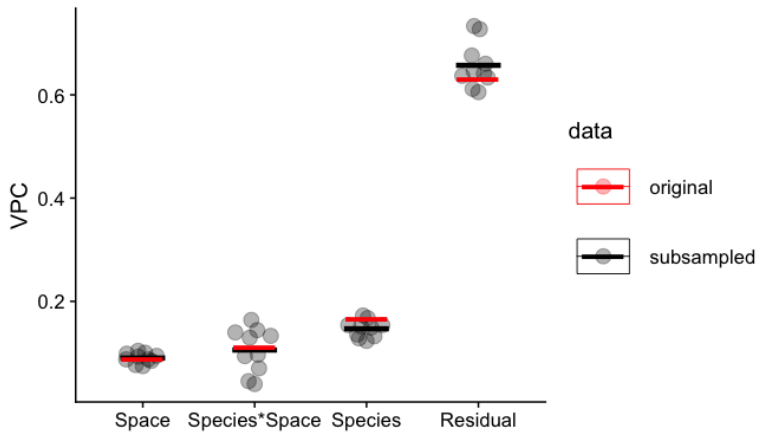


**Figure S2.1**: Comparison of the results for Variance Partition Coefficient (VPC) from growth reduced models (without temporal OPs) at the 5x5 m quadrat scale for Lambir forest plot. Red bars indicate the VPC for the entire 50 ha forest plot dataset and black bars are the mean VPC from 10 subsamples of 5 ha (gray dots).

#### Models with temporal OPs - Fushan and Luquillo forest plots

For the models with temporal OPs, we did the opposite: as the computational time for running models to a huge amount of data are restrictive, we evaluated if the subsampling analysis of 5 ha each plot was able to get the same results if we were doing the analysis with the entire plot. The 5 ha subsamples retained around 85% of the number of species and between 20 and 34% of the number of observations. The vast majority of VPC estimates was very similar between datasets (Figure S2.2). The largest differences (up to 0.03) were found for *species* OP in recruitment for both forest plots, for *time* OP in recruitment for Fushan, and for *species* OP in mortality for Luquillo.

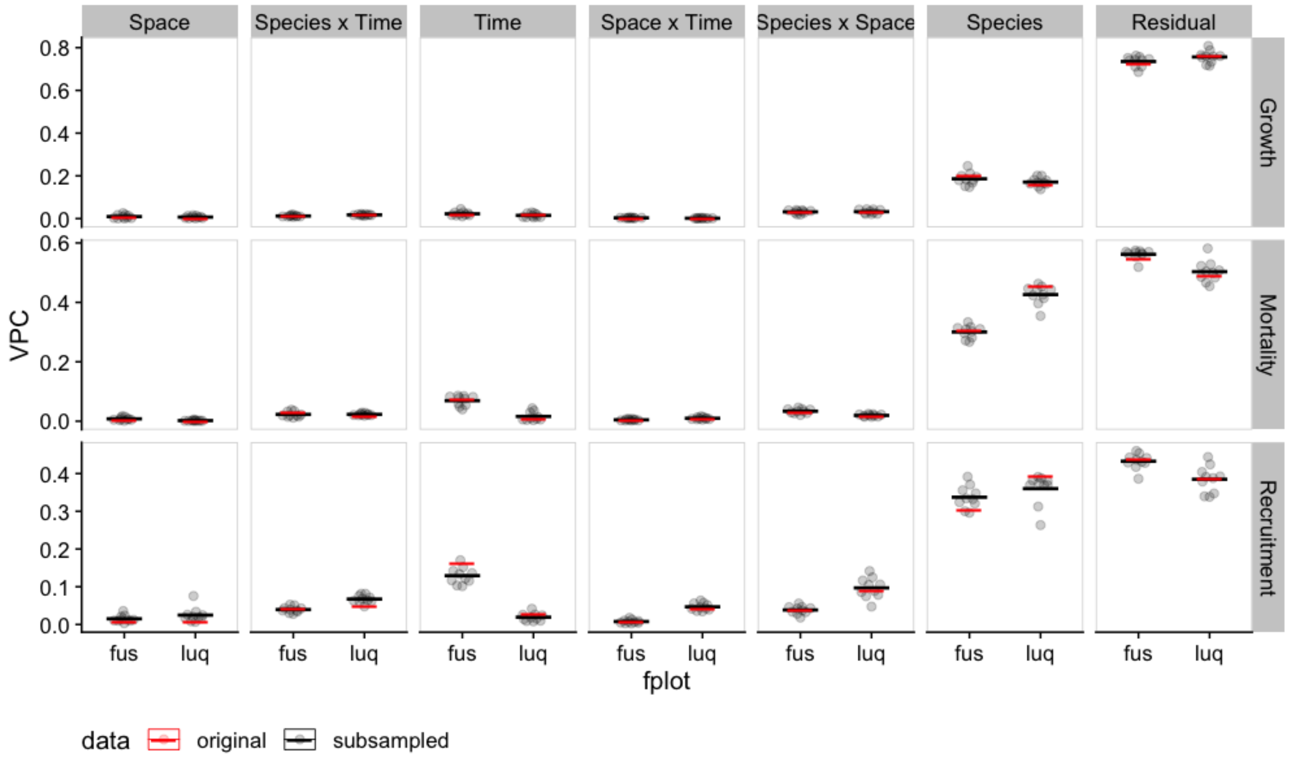


**Figure S2.2**: Comparison of the results for Variance Partition Coeficient (VPC) from the time models at the 100 x 100 m quadrat scale for Fushan (fus) and Luquillo (luq) forest plots. Red bars indicate the VPC for the entire 50 ha forest plot dataset and black bars are the mean VPC from 10 subsamples of 5 ha (gray dots).

### Appendix S3 – Robustness analysis for models with and without temporal organising principles

Given the low number of forest plots with four or more censuses, we could apply the main model of Equation 1 (main text) to only five forest plots (Table S1.1). However, forest plots with just one census interval can also be a reliable source of information for comparing *species*, *space*, and *species x space* OPs. Therefore, we applied a reduced model setup without the temporal OPs - *time*, *species x time*, and *space x time* - for all 21 forest plots. For forest plots with more than one census interval, we averaged the variance estimates across intervals and calculated VPCs.

Using the formula syntax of *brms* R package (Bürkner, 2017), the complete model from equation 1 (hereafter **time models**) is written as:

```
Y ~ 1 + (1|species) + (1|space) + (1|time) + (1|species:space) + (1|species:time) +
(1|space:time)
```

while the reduced model without temporal terms (hereafter **no-time models**) is:

```
Y ~ 1 + (1|species) + (1|space) + (1|species:space)
```

To understand the effects of omitting the temporal terms for the remaining standard deviations and thus the reliability of the reduced model setup, we ran both models for the five forest plots where enough census intervals were available and compared the standard deviations (SD) of the model terms. Below, we show and discuss these comparisons at the 5x5 m quadrat size.

As expected, the total standard deviation in time models was always larger (Figure S3.1) and residual standard deviations for growth (normal distribution) also did not change. Notice that the residual standard deviations for mortality and recruitment are anyway fixed to the theoretical standard deviation of binomial models with complementary log-log link function (Nakagawa *et al.*, 2017). Standard deviations for species organising principle did not change between time and no-time models, except for recruitment where it was slightly larger for some forest plots in no-time models (Figure S3.2). *Space* and *species x space* standard deviations were, in general, smaller in time models for all vital rates, except for growth, where *space* standard deviations were equal or a bit larger for time models (Figure S3.2). It may be the case that *space* and *species x space* OPs in no-time models are incorporating some temporal variability in vital rates, probably through gap dynamics effects on tree recruitment, mortality, and growth. In the lack of temporal OPs, the footprint of fallen-tree gap formation remains in *space* and, at a lower extent, *species x space* OPs.

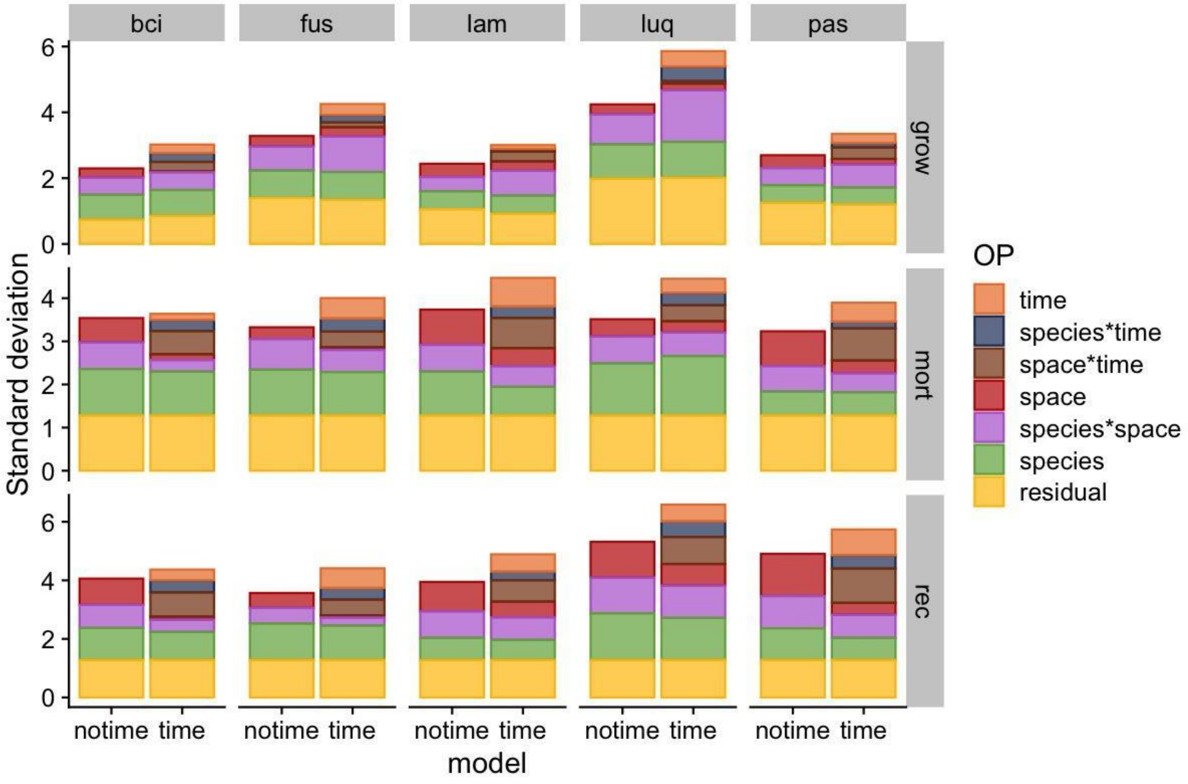


**Figure S3.1.** Standard deviations of models with (time models) and without temporal (no-time models) OPs for the five forest plots with more than four census intervals for growth (grow), mortality (mort), and recruitment (rec) vital rates. Forest plots are Barro Colorado Island (bci), Fushan (fus), Lambir (lam), Luquillo (luq), and Pasoh (pas).

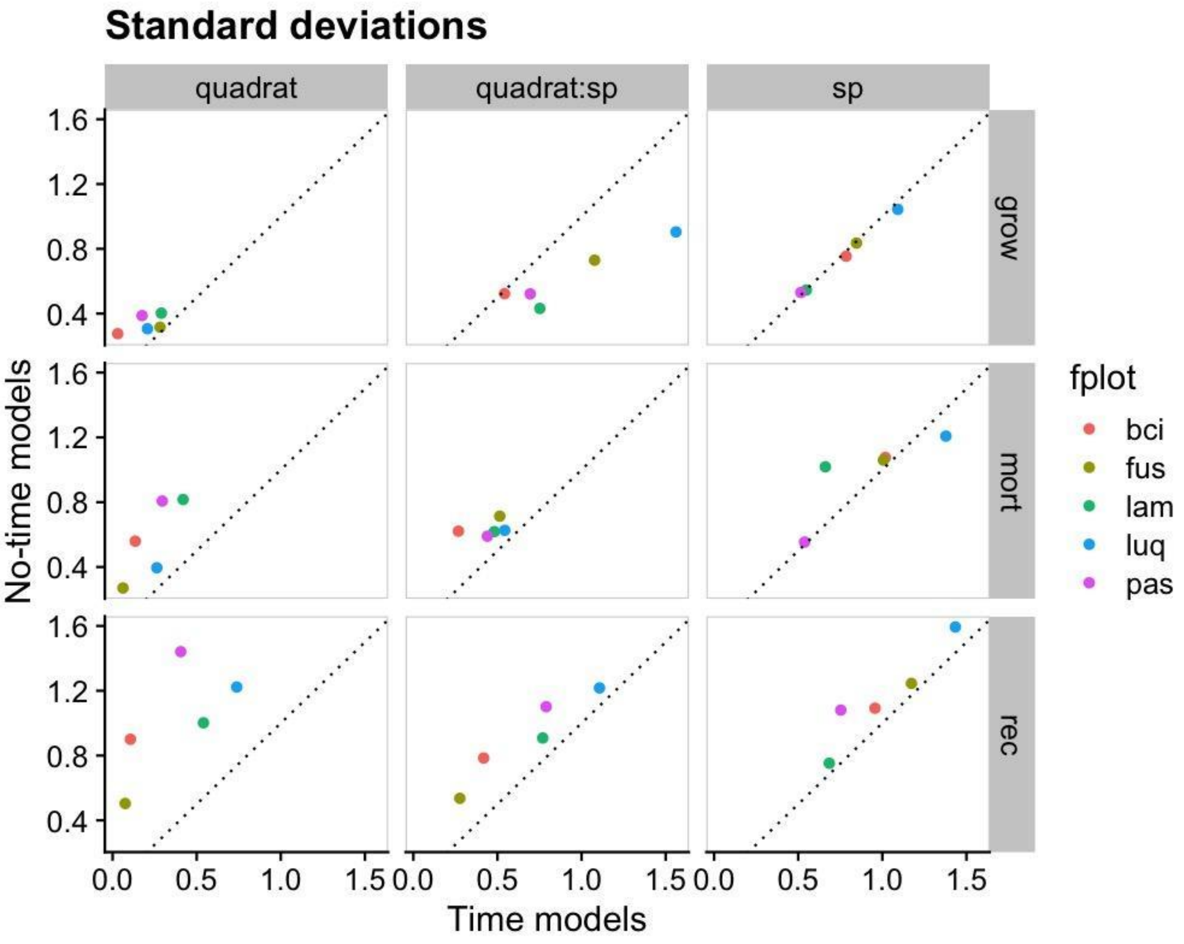


**Figure S3.2.** Comparing standard deviation of *species*, *space*, and *species x space* OPs for models with (time models) and without temporal OPS (no-time models) for the five forest plots with more than four census intervals for growth (grow), mortality (mort), and recruitment (rec) vital rates. Forest plots are: Barro Colorado Island (bci), Fushan (fus), Lambir (lam), Luquillo (luq), and Pasoh (pas). Dotted diagonal lines indicate the 1:1 threshold.

### Appendix S4 - Robustness analysis for the role of rare species

The presence of rare species in a forest plot can influence VPC analyses in the following ways: (1) in multilevel models, rare species may 'shrink' variance estimates towards the population mean because of the small number of observations, and (2) rare species may increase species standard deviations, either by sampling artefact (Condit *et al.*, 2006) or because rare species represent vital rate strategies distinct from the more common species (Umaña *et al.*, 2017). We thus assessed the extent to which rare species affect our results rerunning the VPC analysis (1) without rare species and (2) with rare species grouped as a single 'species'. We used the FuzzyQ clustering algorithm in the 'FuzzyQ' R package (Balbuena *et al.*, 2021) to estimate the probability of each species to be common or rare based on species abundance and occupancy in 50x50 m quadrats. The method has the advantage of allowing comparisons among forest plots of different sizes. Both procedures showed similar results, with a small decrease in the *species* VPC, balanced by an increase in the *residual* and *species x space* VPC when excluding or regrouping rare species. We conclude that our main results are robust to the presence of rare species in the datasets.

#### Classifying rare species

To describe and compare rarity patterns in all forest, we estimated the number of species, number of individuals, and density of rare and common species (Table S4.1). For the forests plots with more than 1 census interval, we averaged the estimates across census intervals. Common species richness ranged from 20% to 56% of total species richness, while it comprised from 86% to 99% of the trees. The average density that formed the cut-off for the rare species classification was 1.39 trees/ha, (Table S4.2).

**Table S4.1.** Number and percentages of species and trees classified as common or rare per forest plot. See Table S1.1 and S1.2 for forest plot names and information. For forest plots with more than 1 census interval, we average the values across intervals.

| | **Species richness** | | | | **Number of trees** | | | |
|---|---|---|---|---|---|---|---|---|
| | **Common** | | **Rare** | | **Common** | | **rare** | |
| **Forest** | **N** | **%** | **N** | **%** | **N** | **%** | **N** | **%** |
| ama | 456 | 35 | 839 | 65 | 98878 | 90 | 11578 | 10 |
| bci | 135 | 42 | 185 | 58 | 334275 | 96 | 12477 | 4 |
| edo | 122 | 29 | 298 | 71 | 168253 | 96 | 7004 | 4 |
| fus | 55 | 50 | 55 | 50 | 143813 | 98 | 2376 | 2 |
| idc | 74 | 53 | 61 | 47 | 34651 | 95 | 1675 | 5 |
| kor | 186 | 40 | 282 | 60 | 335392 | 94 | 22608 | 6 |
| lam | 539 | 38 | 869 | 62 | 376389 | 86 | 61135 | 14 |
| lwd | 14 | 38 | 23 | 62 | 28004 | 96 | 1257 | 4 |
| len | 108 | 29 | 262 | 71 | 145467 | 96 | 4296 | 4 |
| lpl | 108 | 45 | 132 | 55 | 120516 | 96 | 4735 | 4 |
| luq | 55 | 37 | 95 | 63 | 69233 | 96 | 2769 | 4 |
| mos | 94 | 34 | 183 | 66 | 150534 | 95 | 7994 | 5 |
| pas | 375 | 44 | 474 | 56 | 360659 | 92 | 31816 | 8 |
| scbi | 25 | 35 | 47 | 65 | 38366 | 97 | 1248 | 3 |
| serc | 20 | 27 | 55 | 73 | 27031 | 96 | 1205 | 4 |
| sin | 118 | 50 | 118 | 50 | 207356 | 94 | 12196 | 6 |
| ucsc | 10 | 32 | 21 | 68 | 8738 | 96 | 394 | 4 |
| wab | 11 | 28 | 28 | 72 | 49562 | 93 | 3558 | 7 |
| wfdp | 8 | 31 | 18 | 69 | 28948 | 97 | 831 | 3 |
| wyw | 5 | 20 | 20 | 80 | 19458 | 96 | 740 | 4 |
| zof | 3 | 23 | 10 | 77 | 75368 | 99 | 381 | 1 |

**Table S4.2.** Summary values for the density of trees (N/ha) classified as common or rare for the 21 forest plots. SD - standard deviation, Quant - quantiles.

| | **Min** | **Max** | **Mean** | **SD** | **Median** | **Quant 90** | **Quant 95** |
|---|---|---|---|---|---|---|---|
| common | 1.32 | 3840.80 | 35.24 | 128.01 | 11.26 | 63.74 | 109.80 |
| rare | 0.02 | 297.32 | 1.39 | 5.14 | 0.55 | 2.84 | 4.39 |

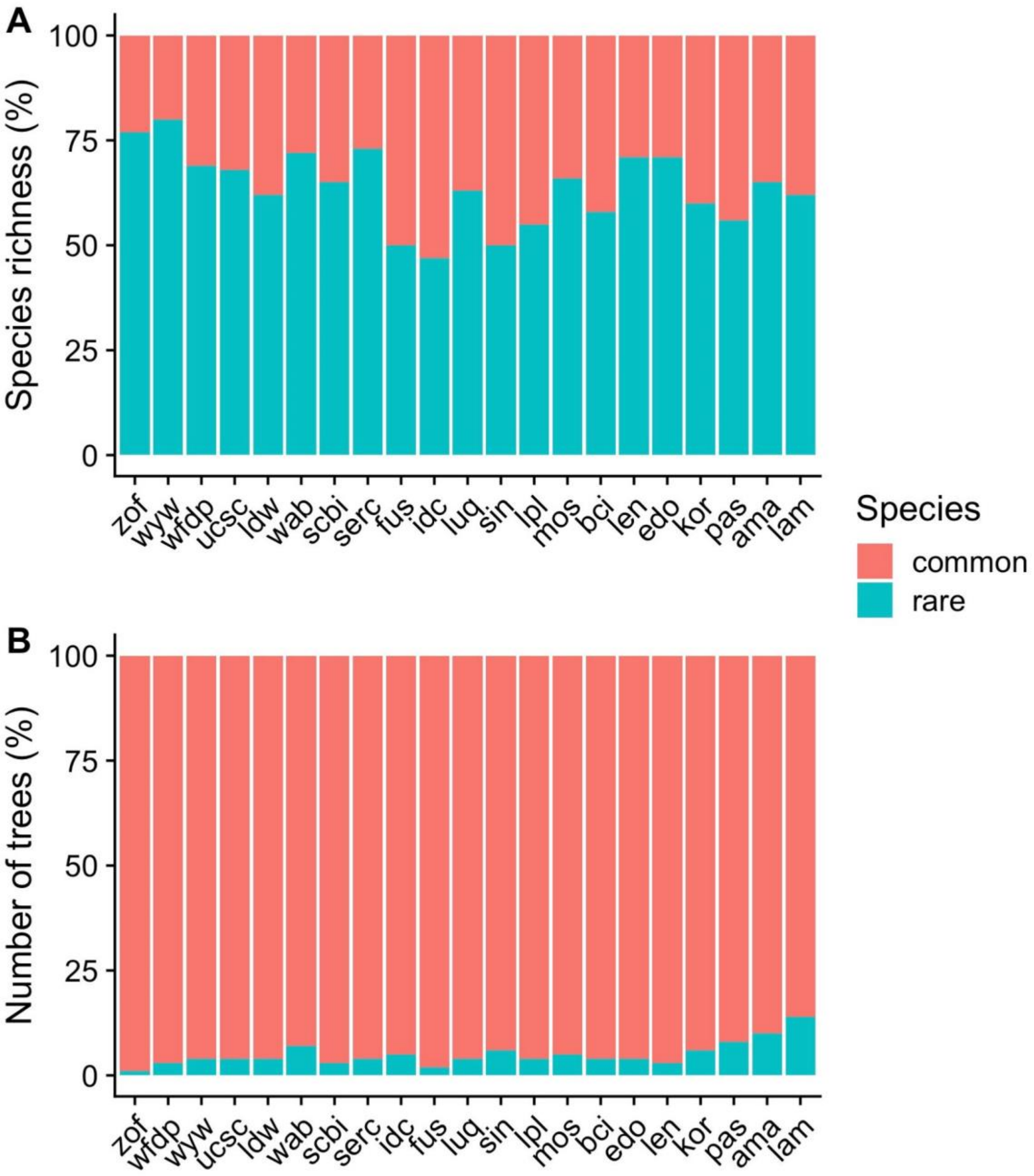


**Figure S4.1.** Percentages of rare and common species and number of trees per forest plot. Forest plots are arranged by absolute latitude (from the largest to the smallest). See Table S1.2 for plots names.

We performed a generalised additive model (Pedersen *et al.*, 2019) to evaluate the relationship between rarity (proportion of rare species) and number of species (at the log10 scale). We found that the percentage of rare species tended to decrease with the number of species only for forests with less than 100 species (Figure S4.2), ranging from 77% in forests with 10 species to 63%

in forests with 90 species. For plots with 100 or more species, the percentage of rare species varied from 62 to 60%.

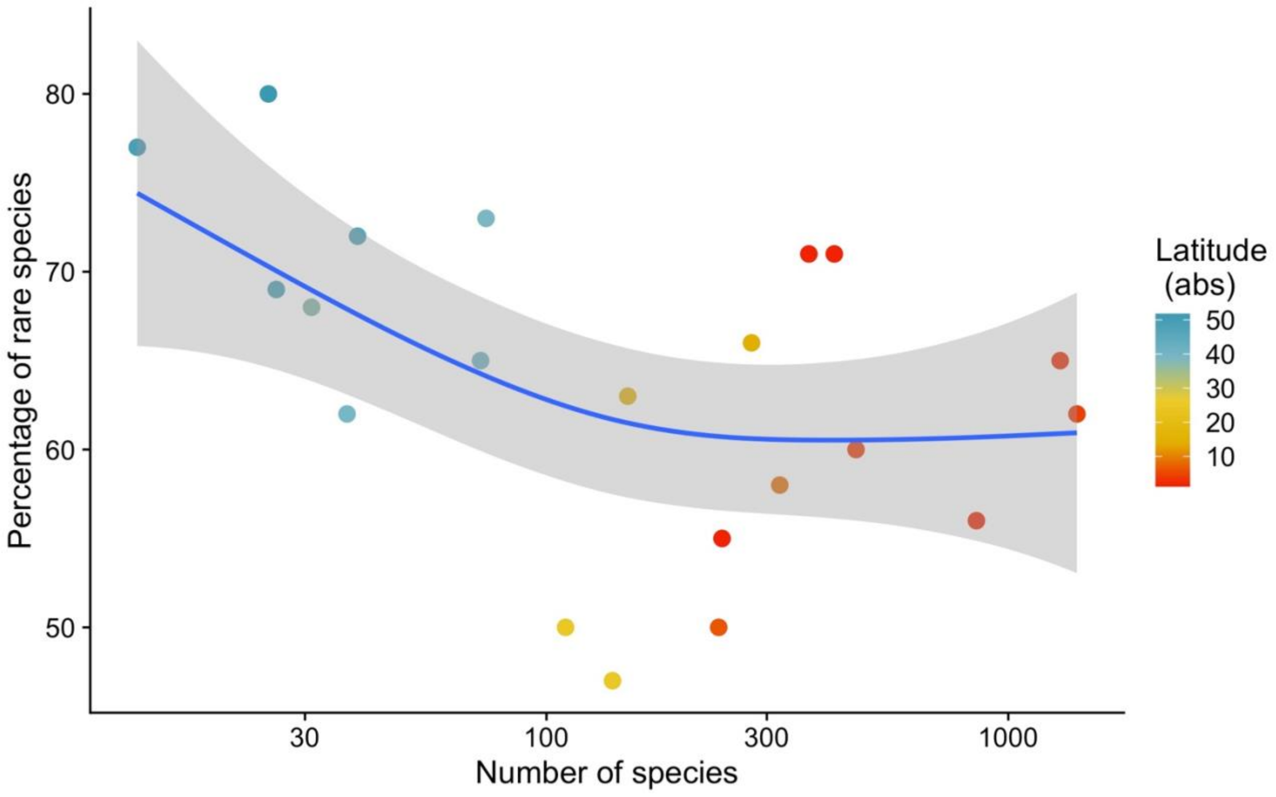


**Figure S4.2.** Percentage of rare species against the number of species (log10 scale) for 21 worldwide distributed forest plots. Blue line and grey area are the fitted results and confidence intervals for a generalised additive model showing a decrease in the percentage of rare species with the number of species but only for forests with less than 100 species. Model's adjusted $R^2 = 0.30$. Each forest plot is coloured by the latitude in absolute values.

#### Excluding or regrouping rare species in forest data

We used two procedures to deal with rare species: (1) excluding rare species from the dataset, which means excluding a proportion of the number of observations in the data, and (2) renaming the rare species the dataset into one generic species name, which does not change the total number of observations in the data. We applied both procedures for the 5 plots with more than 4 censuses using the main model in equation 1 (Fig S4.3) and for all forest plots with the reduced model without temporal organizing principle (Fig S4.4). Both procedures to deal with rare species presented very similar results, and they showed a decrease in *species* VPC, balanced with an increase in the *residual* and *species x space* VPC when excluding or regrouping rare species. For the models with temporal

OPs, we classified the species in the whole dataset as rare and common based on the classification in each census interval (previous section) as some species may temporally vary in abundance/occupancy. We classified a species as rare if it was rare in half or more than half of the time intervals.

The results showed that the largest absolute VPC differences appeared in *species* VPC, decreasing on average from 0.03 (recruitment) to 0.05 (mortality), which corresponds to an average of 11% relative decrease in standard deviation. *Residual* VPC increased on average between 0.01 (recruitment) and 0.06 (mortality). Although *space, time, space x time* and *species x time* standard deviations changed relatively between 3 and 24%, these differences in terms of absolute VPC were negligible (between 0,001 and 0.01). We did not find any tendency for the relative differences in VPC being related to the proportion of rare species in the data (Fig S4.2b).

**Table S4.3:** Average differences in VPC and relative differences in standard deviation between models with rare species and models (1) excluding or (2) regrouping rare species for growth, mortality, and recruitment vital rates. Data used here were the 5 forests with more than 4 census intervals.

| | **Growth** | | | | **Mortality** | | | | **Recruitment** | | | |
|---|---|---|---|---|---|---|---|---|---|---|---|---|
| | **Exclude rare** | | **Regroup rare** | | **Exclude rare** | | **Regroup rare** | | **Exclude rare** | | **Regroup rare** | |
| **Organizing Principle** | **VPC** | **%SD** | **VPC** | **%SD** | **VPC** | **%SD** | **VPC** | **%SD** | **VPC** | **%SD** | **VPC** | **%SD** |
| **space** | 0.002 | 8.8 | 0.002 | 13.3 | 0.001 | 7.9 | 0.003 | 7.9 | 0.001 | 23.3 | 0.002 | 24.3 |
| **species x time** | -0.007 | -13.1 | -0.008 | -23.6 | -0.001 | -5.2 | -0.001 | -6.7 | -0.004 | -5.5 | -0.006 | -10.1 |
| **time** | 0.001 | 4.2 | 0.002 | 10.1 | 0.008 | 3 | 0.008 | 3.3 | 0.008 | 5.3 | 0.011 | 5.4 |
| **space x time** | 0.005 | -4 | 0.001 | -6.9 | 0.008 | 0.6 | 0.002 | -2.9 | 0.004 | 1.1 | 0.013 | 1.8 |
| **species x space** | 0.016 | 2.3 | -0.006 | 0.4 | 0.002 | -0.2 | 0.004 | 2.7 | 0.008 | 2.9 | -0.009 | -3.8 |
| **species** | -0.040 | -11.4 | -0.050 | -12.3 | -0.049 | -12.5 | -0.048 | -12.6 | -0.025 | -8.6 | -0.026 | -10.6 |
| **residual** | 0.023 | 1.4 | 0.059 | 8.2 | 0.030 | - | 0.033 | - | 0.008 | 0 | 0.016 | - |

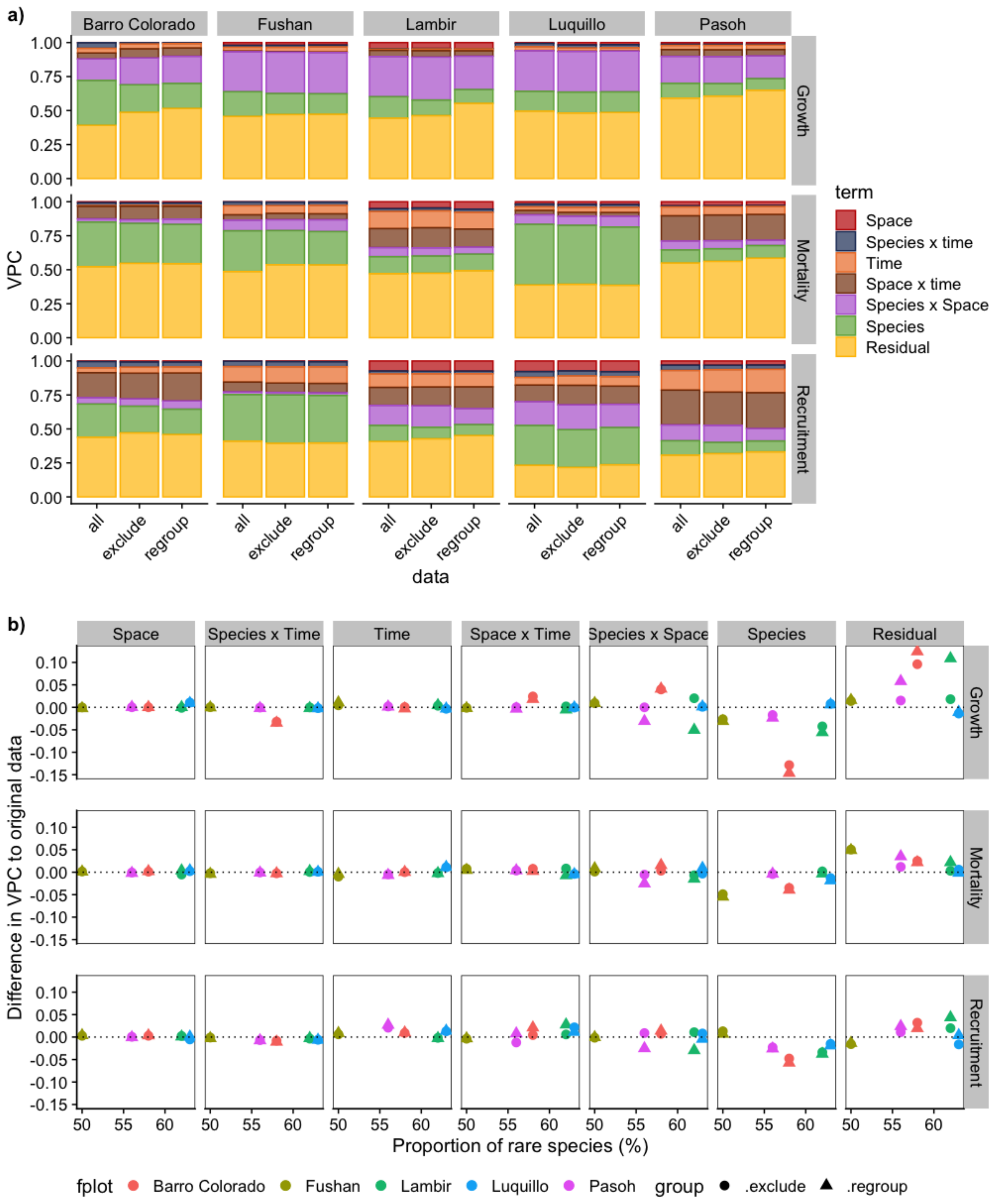


**Figures S4.3. (**a) Comparing Variance Partitioning Components for models with temporal organizing principles among models with all species data included (all), excluding rare (exclude) or regrouping rare species into one generic species label (regroup). (b) Differences in VPC from original data (all species) to the models excluding rare species (circles) or regrouping rare species (triangles). Results here are for the models with the 5x5 m quadrat scale.

For the models without temporal organizing principles (21 forest plots), there was an average of 10 to 15% relative decrease in *species* standard deviation (Table S4.4), while the absolute VPC decreased between 0.03 and 0.09 (Fig S4.3). This decrease was balanced mainly by an increase in *residual* VPCs around 0.02 to 0.07, while for *space* and *space x species* the absolute differences in VPC were very small and on average smaller than 0.01.

**Table S4.4:** Average absolute differences in VPC and relative differences in standar deviations between models with rare species and models excluding or regrouping rare species for growth, mortality and recruitment. Analysis applied to the reduced model without temporal organizing principles for the 21 forest plots.

| | **Growth** | | | | **Mortality** | | | | **Recruitment** | | | |
|---|---|---|---|---|---|---|---|---|---|---|---|---|
| | **Exclude rare** | | **Regroup rare** | | **Exclude rare** | | **Regroup rare** | | **Exclude rare** | | **Regroup rare** | |
| **Organizing Principle** | **VPC** | **%SD** | **VPC** | **%SD** | **VPC** | **%SD** | **VPC** | **%SD** | **VPC** | **%SD** | **VPC** | **%SD** |
| **space** | 0.004 | 11 | 0.004 | 8.7 | 0.008 | 0.0 | 0.006 | -2.6 | 0.006 | -0.6 | 0.013 | 0.5 |
| **species x space** | 0.008 | 8.5 | 0.028 | 4.0 | 0.014 | -1.0 | 0.016 | -0.6 | 0.007 | 0.3 | 0.004 | 0.8 |
| **species** | -0.080 | -11.8 | -0.091 | -12.7 | -0.070 | -14.9 | -0.075 | -16.4 | -0.032 | -9.3 | -0.047 | -10.1 |
| **residual** | 0.070 | 13.2 | 0.084 | 13.9 | 0.048 | 0.0 | 0.053 | 0.0 | 0.017 | 0.0 | 0.019 | 0.0 |

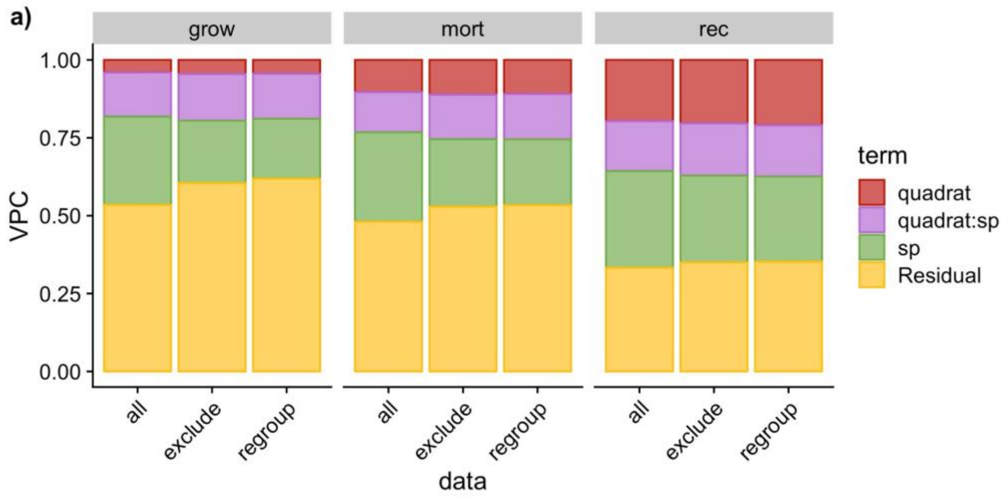

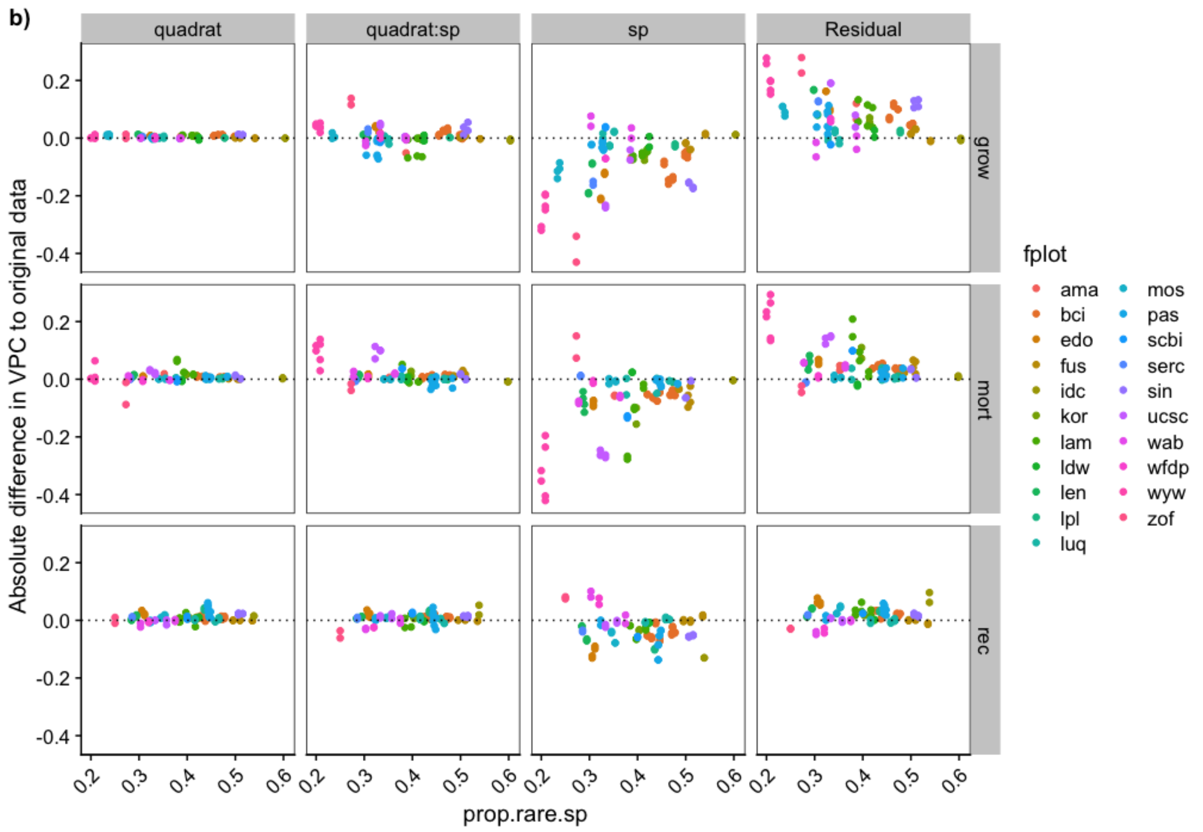


**Figure S4.4.** (a) Comparing Variance Partitioning Components for models without temporal organizing principles (no time models) among models with all species data included (all), excluding rare (exclude) or regrouping rare species into one generic species label (regroup). (b) Differences in VPC from original data (all species) to the models excluding rare species (circles) or regrouping rare species (triangles). Results here are for the models with the 5x5 m quadrat scale.

### Appendix S5 - Additional results comparing global forests

#### Species richness rarefaction

We calculated rarefied species richness based on sampling increment of a quadrat of 20x20 m size. We used the R packages `BiodiversityR` (Kindt & Coe, 2005) and vegan (Oksanen *et al.*, 2020), following Gotelli & Collwel (2001) suggestions for rarefaction curve construction and used the species richness estimated at the smallest plot size (Fig S5.1).

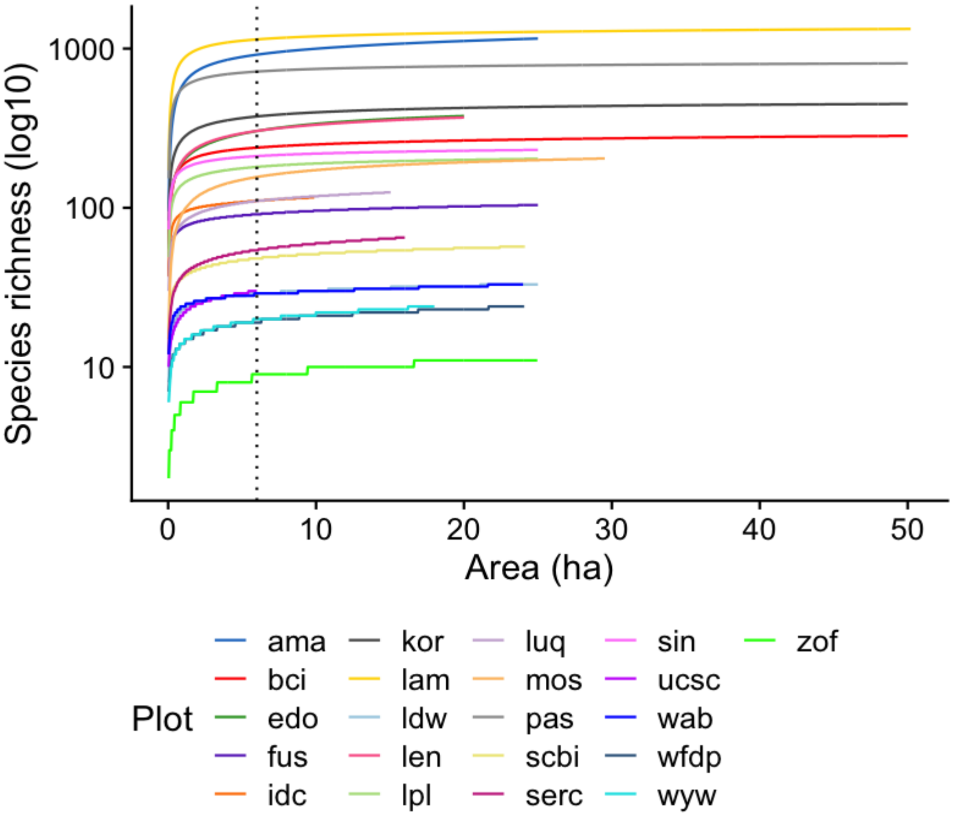


**Figure S5.1.** Rarefaction curves of species accumulations for the 21 forest plots. Vertical dotted line indicates 6 ha area, which is the smallest forest plot area. See Table S1.1 for forest plots abbreviations.

We compare rarefied species richness with other species richness measures, latitude, tree density and metrics of variation in density and richness, with a principal component analysis (Fig. S5.2). All species richness variables were highly correlated and presented the largest contribution to PCA axis 1, which summarised 75.7% of the variation among plots. We, therefore, used the rarefied species richness to compare forest plots.

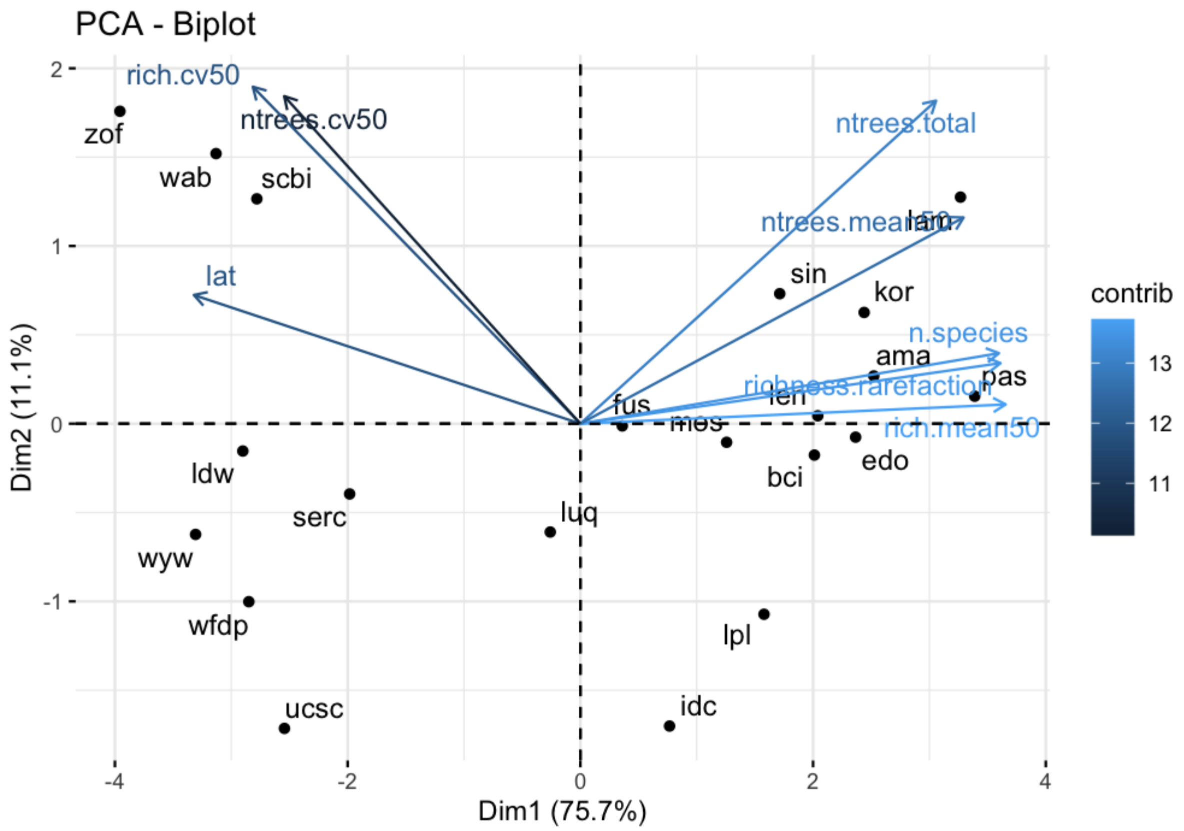


**Figure S5.2**. Principal Component Analysis of the 8 variables used to compare forest plots. All species richness variables were highly correlated and presented the largest contribution to Axis 1 (light blue colours in legend), which summarised 75.7% of the variation among plots. Other variables: latitude (lat), total number of species (n.species), mean number of species (rich.mean50) and coefficient of variation (rich.cv50) in 50 x 50 m quadrat size, total number of trees (ntrees.total), mean number of trees (ntrees.mean50) and coefficient of variation (ntrees.cv50) in 50x50m quadrat size. Richness and tree density variables were log-transformed. See Table S1.1 for forest plots abbreviations.

#### Standard deviation of organising principles across forests

One of the reasons the patterns shown in Figure 4 (main text) - decrease in species VPC with increase in forest species richness - is the increase in standard deviation of other OPs. Here, we investigated how the overall standard deviation of forest vital rates, and each specific organising principle, varies in relation to rarefied species richness. We found that overall standard deviations only decrease for recruitment (FigS5.3) and that decrease is led mainly by the decrease in species standard deviations (Fig. S5.4), which reveals the reason species VPC also decreases with forest species richness.

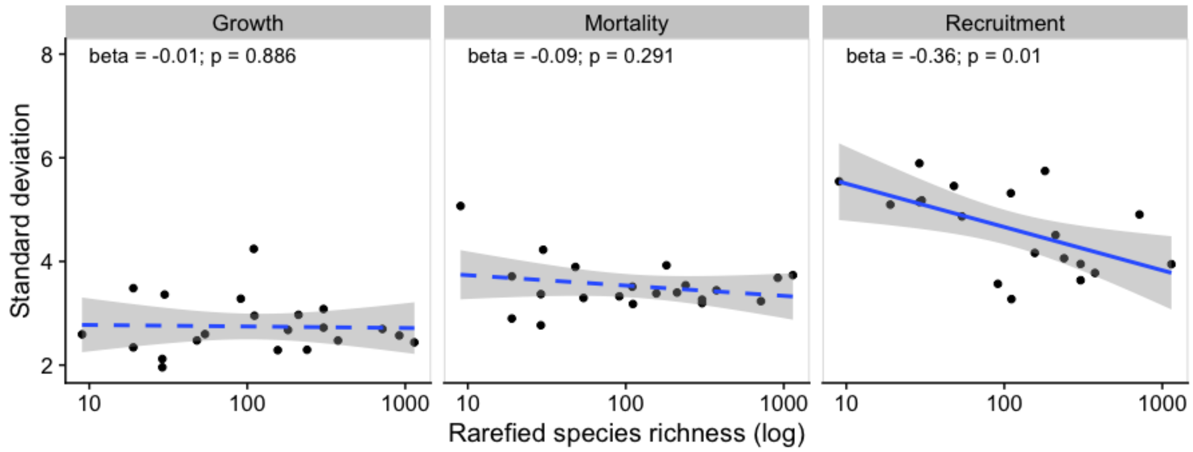


**Fig S5.3.** Overall standard deviation across forest rarefied species richness for the 21 forest plots. Betas (slopes) and their significance were estimated by a linear model with normal distribution.

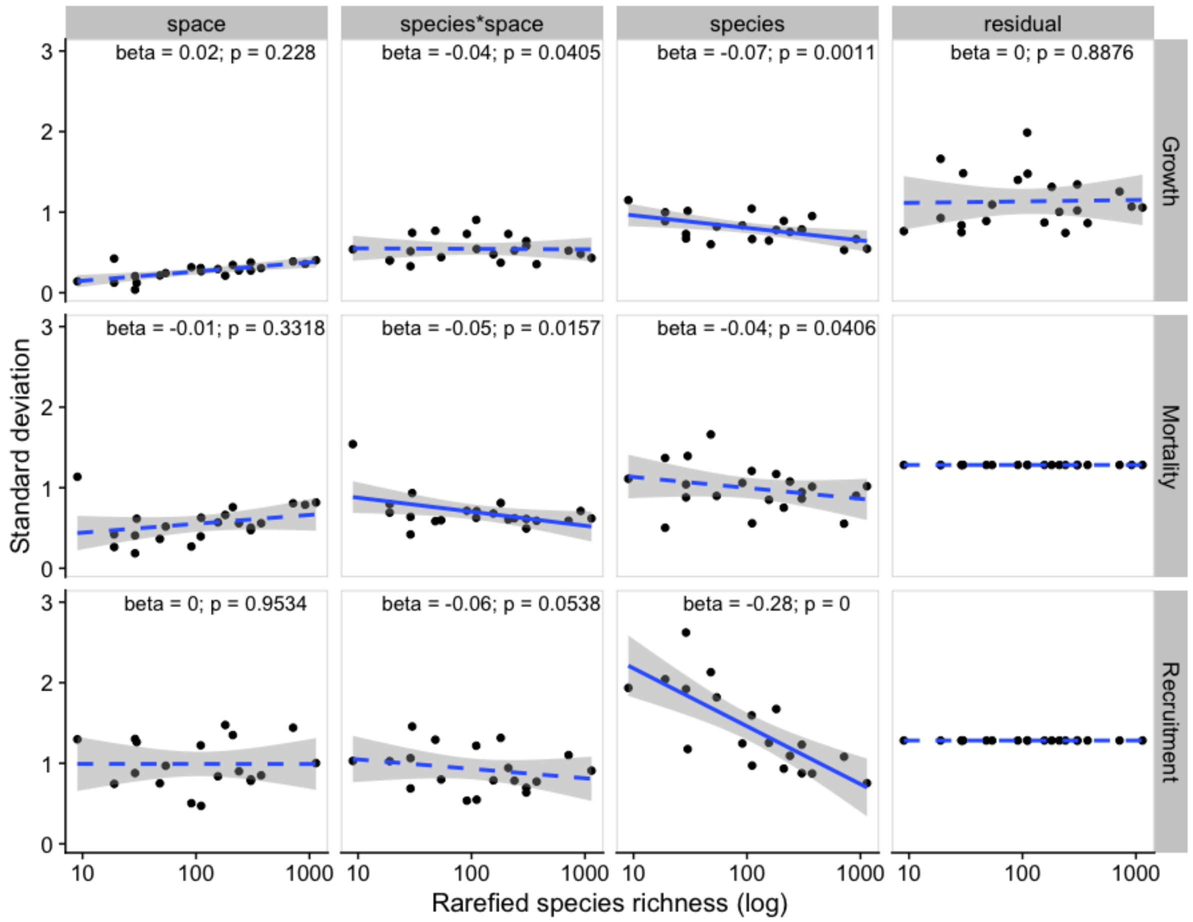


**Fig S5.4.** Standard deviations for each organising principle and vital rates across forest rarefied species richness for the 21 forest plots. Blue dashed lines indicate non-significant betas (slopes) for fitted linear models (normal distributions) and blue solid lines indicate a significant decrease in species standard deviation with increased species richness for recruitment. Multiple tests Bonferroni alpha-level correction was 0.01666.

#### Dirichlet regression models excluding rare species

To test if the results in Figure 4 (main text) are led by differences in rare species richness, we ran the Dirichlet regression with the data excluding rare species. Results were qualitatively similar to the models with all species included Fig S5.5).

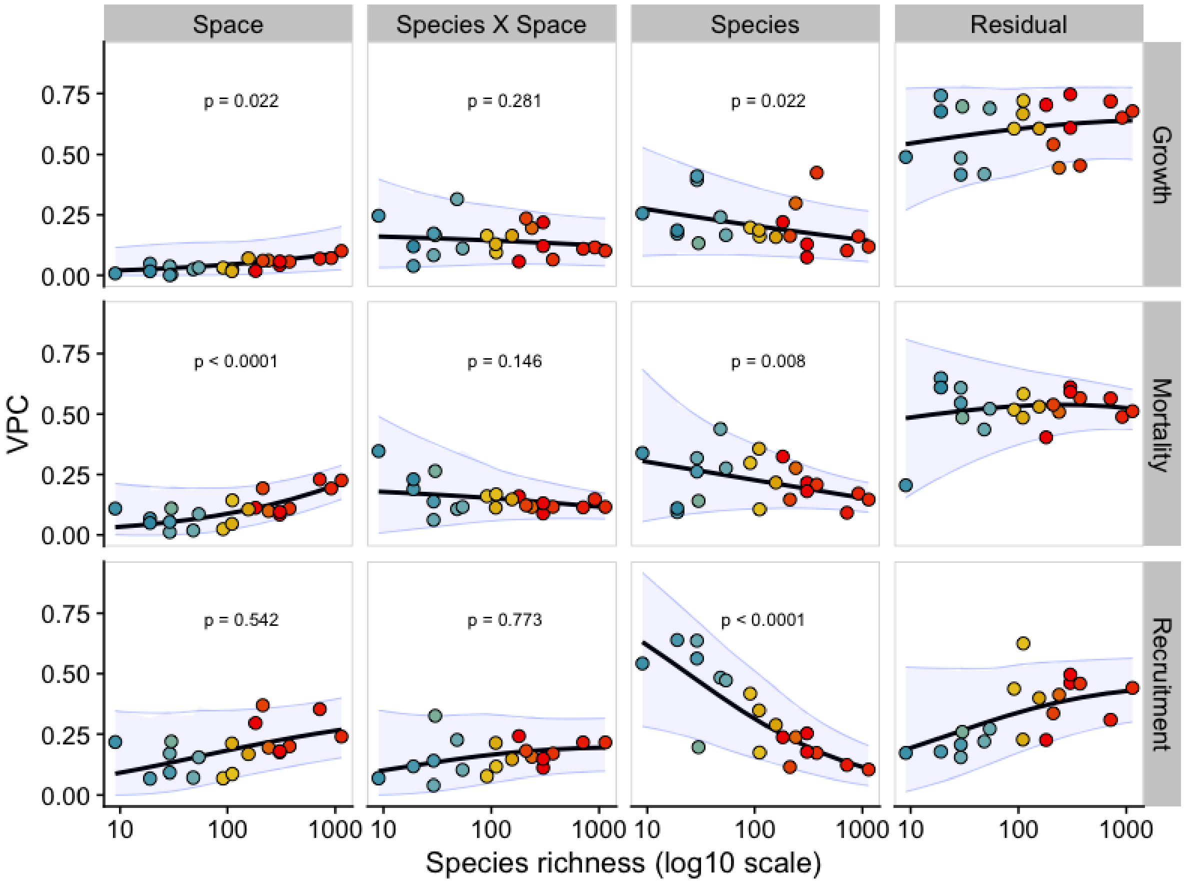


**Figure S5.5**. Dirichlet regression models for the relationship between organising principles VPCs and rarefied species richness applied to forest data excluding rare species. P-values should be compared with alpha after Bonferroni multiple tests correction (alpha=0.016).
